# Altered Socio-Affective Communication and Amygdala Development in mice with Protocadherin10-deficient Interneurons

**DOI:** 10.1101/2024.01.04.574190

**Authors:** Tania Aerts, Anneleen Boonen, Lieve Geenen, Anne Stulens, Luca Masin, Anna Pancho, Annick Francis, Frans Van Roy, Markus Wöhr, Eve Seuntjens

## Abstract

Autism Spectrum Disorder (ASD) is a group of neurodevelopmental conditions associated with deficits in social interaction and communication, together with repetitive behaviors. The cell adhesion molecule Protocadherin10 (*Pcdh10*) has been implicated in the etiology of ASD. *Pcdh10* is expressed in the nervous system during embryonic and early postnatal development and has been linked to neural circuit formation. Here, we show strong expression of *Pcdh10* in the ganglionic eminences and in the basolateral complex of the amygdala at mid and late embryonic stages, respectively. Both inhibitory and excitatory neurons expressed *Pcdh10* in the basolateral complex at perinatal stages and genes linked to vocalization behavior were enriched in *Pcdh10-* expressing neurons in adult mice. To further investigate the involvement of *Pcdh10* in neurodevelopment with relevance to ASD, and to assess the functional and behavioral consequences of loss of Pcdh10 in basolateral amygdala interneurons, we combined a ubiquitous and a conditional Pcdh10 knockout mouse model. Conditional knockout of Pcdh10 reduced the number of interneurons in the basolateral complex. Both models exhibited altered developmental trajectories of socio-affective communication through isolation-induced ultrasonic vocalizations in neonatal pups, characterized by increased emission rates in heterozygous pups. Furthermore, acoustic call features were affected and heterozygous conditional knockout pups emitted calls characterized by reduced peak frequencies but increased frequency modulation. Additionally, we identified distinct clusters of call subtypes with specific developmental trajectories, suggesting the vocalization repertoire is extensive and dynamic during early life. The nuanced alterations in socio-affective communication at the level of call emission rates, acoustic call features, and clustering of call subtypes were primarily seen in heterozygous pups of the conditional knockout and less prominent in the ubiquitous Pcdh10 knockout, suggesting that changes in anxiety levels associated with *Gsh2*-lineage interneurons might drive the observed behavioral effects. Together, this demonstrates that loss of *Pcdh10* specifically in interneurons contributes to behavioral alterations in socio-affective communication with relevance to ASD.

## Introduction

Autism Spectrum Disorder (ASD) is a group of deliberating neurodevelopmental conditions that ranges in both severity and symptoms which can significantly affect quality of life. Individuals with ASD are characterized by deficits in two core domains: persistent deficits in social interaction and communication, together with stereotyped, repetitive patterns of behavior (1). Comorbid disorders include intellectual disability, epileptic seizures, depression, anxiety, fragile X syndrome and attention deficit hyperactivity disorder (2,3). Men seem predominantly affected by ASD, showing a four- to fivefold increase as compared to women (4,5). This is partially caused by the underdiagnosis of high-functioning women in research (6).

Rodent models have proven instrumental translational tools to advance our understanding of genetic and environmental factors involved in ASD etiology. Behavioral phenotyping is a key element of this translational approach and a broad set of sensitive behavioral assays with relevance to the core symptoms and associated features of ASD were developed. These assays target reciprocal social interaction, socio-affective communication, repetitive behavior, sensory processing, motor and cognitive functions and include the assessment of affective changes, such as anxiety, in juvenile and adult rodents (7–10). Considering the significance of early developmental profiles in the disease trajectory of neurodevelopmental disorders such as ASD, it is imperative to study behavioral alterations in young offspring. Because the sensory and motor limitations of neonatal pups complicate the assessment of complex behaviors via standard behavioral assays, isolation-induced ultrasonic vocalizations (USV) are commonly used as the primary indicator for socio-affective communicative impairments at early developmental stages, particularly in rodent models for ASD (8,11,12). Isolation-induced USV are emitted by pups when separated from their mother and littermates, serving important communicative functions to induce maternal retrieval and care (13–15). The whistle-like pup USV span a frequency of approximately 30-110 kHz that peak around postnatal day 6-8 and rapidly decrease thereafter until approximately two weeks after birth (16–18). Anxiolytic drugs, including benzodiazepines and selective serotonin reuptake inhibitors, can decrease the emission rate and duration of these vocalizations, suggesting the mitigation of a presumed anxious state (19–23).

*Protocadherin10 (Pcdh10)* is a cell adhesion molecule of the δ2-subfamily of protocadherins (24). Protocadherins are believed to mediate intracellular signaling via their unique cytoplasmic tails, in conjunction with their weak adhesive properties (25). Mutations or copy number variations in *PCDH10* have been linked to ASD (26,27) and comorbid disorders such as Tourette’s syndrome in humans (28). *Pcdh10* is predominantly expressed in the nervous system, including neurons of the olfactory system, limbic system, visual system, cerebellum and spinal cord (24,29–32). During early embryonic development, *Pcdh10* expression is most prevalent in the subpallium, increasing in the cortex starting from embryonic day (E)13.5 (33). Deletion of *Pcdh10* in mice, by replacing the first exon with a LacZ-neo cassette, resulted in postnatal lethality and severe axonal defects (33). A recent study using a *Pcdh10* knockout (KO) line without insertion of a *LacZ-neo* cassette refuted these findings (34). Despite the disparities, *Pcdh10* deficiency was associated with altered behaviors reminiscent of the core symptoms of ASD in both mouse models. This included decreased social approach and alterations in socio-affective communication through USV, together with changes in anxiety levels, impaired fear learning and reduced stress coping (34–36).

Perturbed balance between excitatory neurons and inhibitory interneurons is widely believed to play a prominent role in ASD pathophysiology (37–40). Interneurons deliver the necessary inhibitory input into local circuits, synchronize projection neuron populations and stabilize network dynamics (41). Multiple ASD mouse models show a reduced amount or signal intensity of interneurons in the cortex (42–46), hippocampus (43,44), striatum (42,44) and amygdala (47). One of the hallmarks of ASD mediated by interneurons includes reduced gamma band oscillations (48). Heterozygous *Pcdh10*^+/LacZ^ mice show reduced power of gamma band activity from the lateral (LA) to basolateral (BLA) amygdala in *ex vivo* brain slices (36) and of gamma band auditory steady state responses (49). The high expression of *Pcdh10* in interneuron-rich subpallial structures during development and its association with gamma band oscillations imply that the effects of reduced *Pcdh10* might be caused by an excitation/inhibition imbalance.

In this study, we investigated the role of *Protocadherin10* in altered neurodevelopment with relevance to the etiology of ASD using two mouse models. The first model has a full body deletion of *Pcdh10*, while the second model has only selective deletion of *Pcdh10* in *Gsh2-*lineage interneurons. We found that *Protocadherin10* expression in subpallial interneurons increased from embryonic to early postnatal life and was particularly strong in the basolateral amygdala. Conditional knockout of *Pcdh10* in *Gsh2-*lineage interneurons reduced this population in the basolateral amygdala. Strikingly, heterozygous loss of *Pcdh10* in the conditional model paralleled, and surpassed, the behavioral phenotype seen in the full knockout, marked by an increase in isolation-induced USV emission rates, decreased peak frequency and increased frequency modulation. Additionally, we identified distinct clusters of call subtypes and observed nuanced changes between the genotypes of the conditional knockout model. Alterations in socio-affective communication in the conditional knockout suggest *Gsh2*-lineage interneurons drive the observed behavioral effects, and loss of *Pcdh10* specifically in interneurons contributes to behavioral alterations relevant to ASD.

## Methods

### Animals

Mice were generated from a mutant line containing a floxed first exon of the *Pcdh10* gene (50), backcrossed in a CD-1/Swiss background for at least nine generations. Pcdh10^fl/+^ mice were crossed with either a RCE line (51) or a Gsh2-iCre line (52). In order to label and selectively delete *Pcdh10* expression in *Gsh2-*interneurons, female Pcdh10^fl/fl^;RCE^fl/fl^ mice were mated with male Gsh2-iCre^+/-^;Pcdh10^fl/+^ mice. Homozygous, Cre-positive offspring (Gsh2-iCre^+/-^; Pcdh10^fl/fl^; RCE^fl/+^) are referred to as ‘Gsh2:cKO’ mice, while their heterozygous, Cre-positive littermates (Gsh2-iCre^+/-^; Pcdh10^fl/+^; RCE^fl/+^) were used as ‘Control’. A breeding scheme is shown in Supplementary Figure 1. To generate the ubiquitous *Pcdh10* knockout (KO) mice, male Pcdh10^fl/+^ mice were crossed with female *Pgk1*-Cre^+/-^ mice (53) and Pcdh10^+/-^ offspring was backcrossed in a CD-1 background for at least three generations. For timed experiments on embryos, mice were mated and females were checked for a copulation plug every morning. This day was considered E0.5. Mice were housed at KU Leuven in a conventional facility with a 14/10 light/dark cycle in a humidity- and temperature-controlled room.

### Genotyping

Polymerase Chain Reaction (PCR) was performed on purified DNA originating from ear or tail biopsies. Specific amplicons were prepared using several primer pairs (Table 1) in combination with GoTaq© DNA Polymerase (Promega).

**Table 1:**
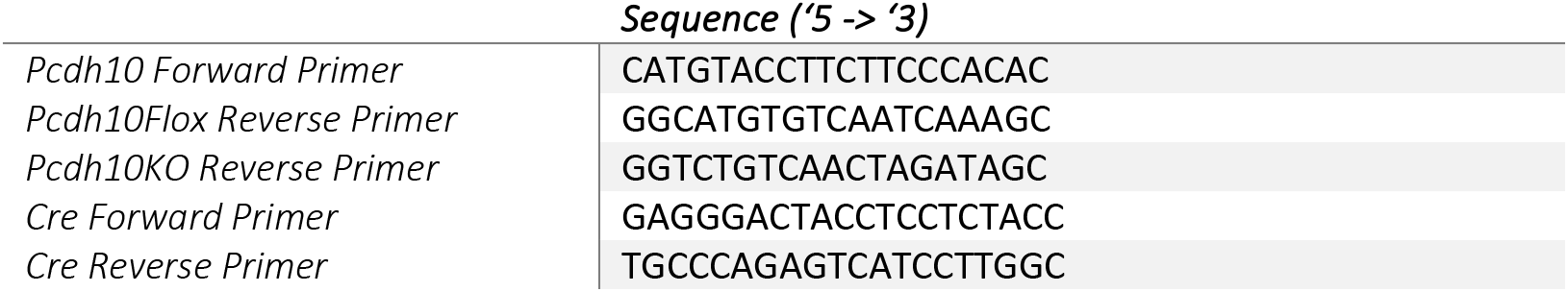
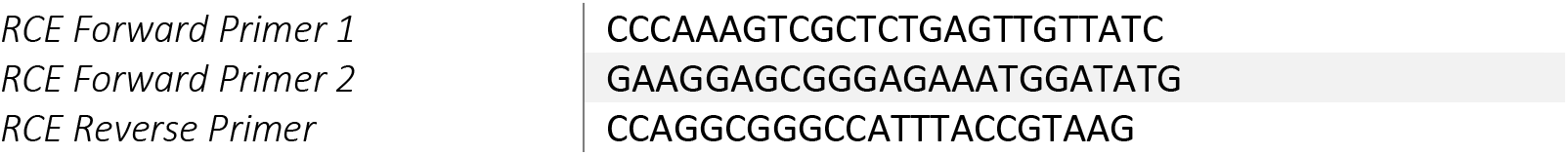
Genotyping primers for Pcdh(c)KO mice.

### Tissue processing

Collected postmortem brain samples were isolated in ice-cold phosphate-buffered saline (PBS, 8 mM Na_2_HPO_4_.2H_2_O, 2 mM KH_2_PO_4_, 150mM NaCl, 3 mM KCl, pH = 7.4), followed by overnight fixation in 4% paraformaldehyde (PFA) in PBS at 4°C. Juvenile pups used for postmortem brain isolation were anesthetized with an intraperitoneal injection of 50µL dolethal (Sodiumpentobarbital, 2mg/kg), followed by intracardiac perfusion with saline and 4% PFA. After overnight fixation, samples were washed with PBS and stored in storage buffer (PBS with 0.4% NaN_3_) at 4°C until use. Brains or tissue used for cell culture or flow cytometry were collected in ice-cold L15^++^ media (Leibovitz’s L-15 media (Gibco) supplemented with 35 mM D-glucose (Sigma) and 2,5 mM HEPES (Gibco)). For protein isolation, brains were collected in ice-cold PBS supplemented with cOmplete, EDTA-free Protease Inhibitor Cocktail Tablets (Merck) and snap frozen in liquid nitrogen before storage at −80°C. Ganglionic eminence tissue used for qPCR analysis was isolated in ice-cold PBS and stored at - 20°C until use.

### Paraffin embedding

Fixed brain tissue was placed in cassettes, washed overnight with saline, and then subjected to an alcohol dehydration protocol and overnight paraffin embedding protocol with the Excelsior™ AS Tissue Processor and the HistoStar™ Embedding Workstation. Embedded tissue was placed on a wooden holder and inserted into the Microm HM360 rotary microtome. Coronal sections of 6µm were collected on SuperFrost® Plus slides (Thermo Fisher Scientific) and dried overnight. Sections were kept at room temperature until use.

### Vibratome sectioning

Fixed brains were embedded in 4% ultrapure agarose (Thermo Fisher Scientific) and submerged in cutting buffer (8 mM Na_2_HPO_4_.2H_2_O, 2 mM KH_2_PO_4_, pH = 7.4) in the Micron HV650 (Thermo Fisher Scientific). Coronal sections of 60µm were cut using a frequency of 70Hz, an amplitude of 0.7mm and a speed of 1.4mm/s. Sections were collected in storage buffer and stored at 4°C until use. To analyse eGFP signal in Gsh2:cKO and Gsh2:Control pups, vibratome sections were counterstained with 1:1000 DAPI (4ʹ,6-diamidino-2-phenylindole, 1 mg/ml, Sigma) in PBS and incubated for 20 minutes in the dark, before mounting with Mowiol (Sigma).

For Hybridization Chain Reaction (HCR) on floating vibratome sections, vibratome sections were cut in a RNAse free environment; storage buffer and cutting buffer were prepared in autoclaved PBS-DEPC (Acros), agarose was prepared in UltraPure DNase/RNase-Free Distilled Water (Invitrogen) and razor blades and cutting bath were cleaned with RNase AWAY^TM^ Surface Decontaminant (Thermo Fisher Scientific). Sections were collected as described and used for HCR procedures within the week.

### *In Situ* Hybridization

10µg of a plasmid containing 807bp of *Pcdh10* exon 1 sequence was linearized with SacII (NEB) in 1x CutSmart buffer (NEB). *In vitro* transcription was performed by adding linearized plasmid, Sp6 polymerase (Roche), DIG labeling mix (Roche) and RNase inhibitor (Roche) in 1x reaction buffer (Roche) and incubating overnight at 37°C. Generated probes were purified using Micro Bio-SpinTM P-30 Gel Columns with RNase-free Tris Buffer (BioRad) according to the manufacturer’s protocol and stored at −80°C. *In Situ Hybridization* (ISH) was performed using an automated system (VENTANA Discovery, Roche). Paraffin sections were first processed in a series of deparaffination, post-fixation, pretreatment, and proteinase K digestion steps. 200ng of antisense *Pcdh10* probe in Ribohybe (Roche) was added to the sections, followed by denaturation (6min, 90°C) and hybridization (6hrs, 70°C). Sections were washed with 0.1X SSC buffer (Roche) three times at 68°C for 12 minutes each, and post-fixed in RiboFix (Roche). An anti-DIG-alkaline antibody (Roche) was added to the sections, followed by blocking, addition of the BCIP/NBT substrate (BlueMap1 and BlueMap2) and incubation for 7 hours to form a blue precipitate. Sections were washed, counterstained using the Red Counterstain II kit (4 min, Roche), removed from the automated system and rinsed with distilled water. This was followed by progressive ethanol and xylol dehydration and mounting (Eukitt quick-hardening mounting medium, Sigma).

### Hybridization Chain Reaction

Hybridization chain reaction (HCR) probe pairs targeting the gene of interest were designed using the Jupyter Notebook application in Anaconda with an optimized script as described in (Elagoz et al., 2022). Probes were ordered as 50pmol DNA oligopools from Integrated DNA Technologies (IDT), dissolved in 100µL nuclease-free water, aliquoted, and stored at −20°C until use. The used protocol was based on the HCR v3.0 protocol published by Choi et al. (2018) (54). Paraffin sections were processed via progressive deparaffinization, rehydration and permeabilization steps (1:3000 Proteinase K (Roche) in PBS-DEPC, 5min, 37°C). Sections were then washed in autoclaved milliQ, prehybridized by adding probe hybridization buffer to the sections and incubating (30min, 37°C). 1,2pmol probe mixture was prepared in probe hybridization buffer per slide, and incubated for at least 16 hours at 37°C. Following incubation, excess probe was removed by progressive washing in a series of increasing 5x SSCT buffer (5x SSC, 0,1% Tween20) in probe wash buffer for 15 minutes each at 37°C. Sections were pre-amplified by adding amplification buffer and incubated (30min, RT). 9pmol of the appropriate hairpin 1 and hairpin2 were separately heated at 95°C for 1,5 minutes per section, followed by cooling on ice for 5 minutes and incubation in the dark (30min, RT) before adding them to the sections. Sections were incubated at room temperature in the dark for at least 16 hours. To remove excess hairpins, sections underwent three consecutive washes with 5x SSCT buffer for 10 minutes each. Sections were counterstained with 1:1000 DAPI in 5x SSCT and incubated for 20 minutes in the dark. A final wash with 5x SSCT buffer was done before mounting with Mowiol (Sigma) and storage at 4°C until imaging with Olympus FluoView FV100.

For HCR on vibratome sections, sections were permeabilized with a solution of 0.5% Triton-X100 in autoclaved PBS for two hours at 37°C in the dark. Sections then underwent prehybridization, probe hybridization, washing, pre-amplification, amplification and DAPI staining with solutions and time schedule as described, however probe concentration was increased 3,33 times, incubated for at least 20 hours and sections were processed as floating sections on a shaking table instead.

### Imaging

All imaging of fluorescent sections was done using the Olympus FluoView 1000D (confocal) or Leica DM6 (epifluorescence) microscope. Images of *in situ hybridization* (ISH) were taken with the Leica DM6 in brightfield mode. Images were processed with Fiji/ImageJ (55). To calculate the percentage of eGFP^+^ interneurons in the basolateral complex (BLC), a max projection based on intensity values was made. The BLA/LA outline was traced and a local threshold (Bernsen) was applied to the green channel. Threshold was then converted into a mask and area was measured.

### Flow cytometry and Fluorescent Activated Cell Sorting (FACS)

Gsh2:Control and Gsh2:cKO embryos were identified by their presence of eGFP signal when observed under a SteREO Discovery.V8 binocular fluorescence microscope (Zeiss). For each experiment, appropriate negative and single-stained controls were taken along. The telencephalon or ganglionic eminences were isolated, dissociated with forceps and digested at 37°C for 30 minutes via the addition of digestion solution (6 units of Papain (Sigma) and 9 units of DNase I (Sigma) in L15^++^ media). Cells were washed three times with ice-cold DPBS (Gibco). Hereafter, cells were suspended in 100µL DPBS and 10µL 7-AAD (BD Biosciences) was added and incubated for 15 minutes at room temperature. 400µL DPBS was added and cells were strained over a Falcon round-bottom tube with a 35µm Cell Strainer Cap (Corning). Cells were analysed or sorted using the SH800S Cell Sorter equipped with SH800S software (Sony). To analyse flow cytometry data, FlowJo (Becton, Dickinson & Company) or FCS Express 7 (De Novo Software) applications were used. For sorting, a 100µM sorting chip (Sony) was used and cells were sorted using the semi-purity or purity settings. Approximately 300000 cells were collected in RNAse-free Microfuge tubes (Thermo Fisher Scientific) containing 500µL ice-cold DPBS and 1µL Protector RNAse inhibitor (Roche). For subsequent analysis of mRNA expression via qPCR or bulk RNA sequencing, cells were spun down (1000g, 5min), resuspended in 1mL TRIzol™ Reagent (Life Technologies), vortexed for 60 seconds and incubated at room temperature for 5 minutes. Hereafter, cells were snap-frozen in liquid nitrogen and stored at −80°C. For analysis of DNA content via qPCR, cells were spun down (1000g, 5min), suspended in 200µL nuclease-free water and frozen at −20°C.

### Quantitative Polymerase Chain Reaction (qPCR)

Total RNA content was extracted from E13.5 ganglionic eminence tissue via the RNeasy™ Plus Mini Kit (Qiagen). Briefly, tissues were thawed, 1mL TRIzol reagent (Invitrogen) was added and tissue was homogenized using a drill. After a brief incubation with 200µL chloroform, a density centrifugation at 12000g (3 min) was performed and the upper aqueous phase with RNA was collected and washed with an equal volume of 70% ethanol. RNA was then extracted via the RNeasy MinElute spin column following the manufacturers protocol. The RNA content was eluted in 50 µl nuclease-free water and stored at −80°C if not used immediately for cDNA synthesis. For qPCR on sorted cells, cells frozen in Trizol were thawed on ice and RNA was extracted with the RNeasy™ Plus Micro Kit (Qiagen) and eluted in 14µL nuclease-free water. Separate biological samples with the same genotypes were sometimes pooled together in order to increase RNA yield. For qPCR starting from DNA, DNA was purified via isopropanol/ethanol precipitation and purified DNA was dissolved in 50µL nuclease-free water.

cDNA was synthesized using the SuperScript® III Reverse transcriptase kit according to the manufacturer’s protocol (Invitrogen). 2 units of Ribonuclease H (Invitrogen) were added at the end of the procedure to remove any remaining RNA. A non-template control (NTC) and non-reverse transcriptase control (NRTC) were included in every experiment. qPCR reactions were set up using the SYBR Green supermix containing the SsoAdvanced^TM^ Polymerase (BioRad), 500nM forward and reverse primer and 1/1-40 diluted cDNA sample. A CFX96 Real-Time System (BioRad) was used for thermocycling. The used primer pairs are shown in Table 2. Primers targeting housekeeping genes GAPDH and B-actin were used for normalization of gene expression. Relative, normalized gene expression was calculated using the CFX Maestro software.

**Table 2:**
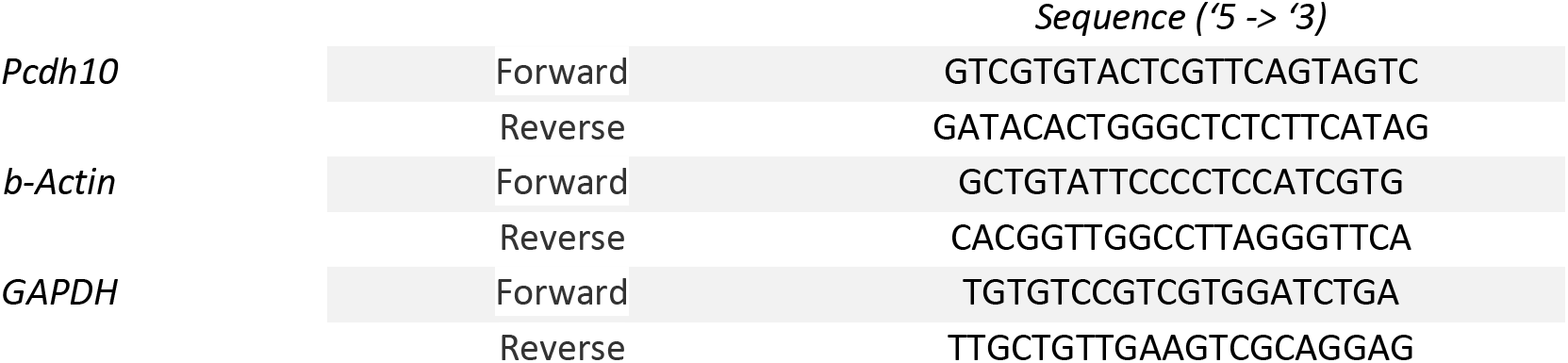
qPCR primers used.

### Bulk RNA Sequencing

RNA samples of sorted *Gsh2-*lineage cells (eGFP^+^) from control and cKO embryos (E15.5) were submitted for quality control with the BioAnalyzer (Agilent) at the Genomics Core (KU Leuven). 12 samples were selected for library preparation and sequencing. Libraries were prepared using the QuantSeq 3’ mRNA-Seq Library Prep Kit FWD (Lexogen) and samples were sequenced on the HiSeq4000 (Illumina) in single read mode at the Genomics Core (KU Leuven). FastQC (56), accessed via the Galaxy webserver (usegalaxy.eu), was used for initial quality check of raw reads. Subsequent data analysis was done using the Flemish Supercomputer Center. Adapter contamination, polyA and low quality tails were trimmed using BBDuk (57) in combination with Lexogen i7 and i5 Index Sequences. Indices for genome alignment were generated from the *Mus musculus* GRCm39 genome and GRCm39.108 gene files, available at ensembl.org/Mus_musculus/Info/Index. Reads were aligned with STAR aligner (58) and Featurecounts (59) was used to quantify the reads. Hereafter, reads were analysed using DESeq2 (60).

### Western Blotting

Snap-frozen samples were lysed with ASBA lysis buffer (61) containing protease inhibitor cocktail solution (Roche), followed by mechanical homogenization (5x 10sec), sonification (5x 10sec), incubation (5min, 70°C) and centrifugation (15min, 13000rpm, 4°C). Supernatant was collected and stored at −80°C. Protein concentration was determined using the Qubit Protein Assay Kit (Invitrogen) and Qubit Instrument (Thermo Fisher Scientific) according to the manufacturer’s protocol.

For Western blotting, 20µg sample was mixed with sample buffer (BioRad) and reducing agent (BioRad). Samples were denatured for 10 minutes at 70°C and proteins were separated (55min, 200V) on a Midi 4-12% Bis-Tris gel (BioRad) in 1x MOPS buffer (BioRad) along with 4µL of Spectra multicolor Broad Range marker (Thermo Fisher Scientific). Proteins were transferred to a Polyvinylidene difluoride membrane (PVDF, BioRad) using the Trans Turbo Blot (BioRad) for 15 minutes. The PVDF membrane was rehydrated in Tris buffer (5min, 10mM Tris, 100µM thimerosal, 150 mM NaCl, 0,01% Triton-X100, pH=7.6), and incubated in Blocking Buffer containing 5 % Blocking for ECL (GE healthcare) in Tris buffer for two hours. Rat anti-PCDH10 antibody (Anti-OL-protocadherin antibody, clone 5G10, Sigma) or mouse anti-GAPDH (Merck) was diluted 1:1000 in blocking buffer, added to the membrane and incubated overnight. Blots were rinsed three consecutive times with Tris buffer (5min) and 1:10000 diluted donkey- or goat-anti-rat-HRP (Agilent Technologies) or 1:25000 diluted goat-anti-mouse-HRP (BioRad) in blocking buffer was added (30min). The blot was subsequently washed with Tris buffer (2x 5min) and Tris-stock (1x 5min, 50mM Tris, pH = 7.6) and incubated for 5 minutes in ECL solution for chemiluminescent detection (Thermo Fisher Scientific). Proteins were visualized using the BioRad Imaging system with Chemi visualization.

### Ultrasonic Vocalizations

Female mice were mated with appropriate stud males and individually housed upon visible signs of pregnancy. Hereafter, cages were inspected daily for signs of birth. Isolation-induced pup USV were recorded on postnatal days P3, 6, 9 and 12 during the light cycle at room temperature. The number of pups used per genotype and age is shown in Supplementary Table 1. To induce pup ultrasonic calling, pups were taken from their home cage and immediately placed into a glass container with fresh bedding positioned in a soundproof styrofoam box, applying a protocol previously established in the laboratory (16). USV were recorded for a period of 10 minutes using the UltraSoundGate Condenser Microphone CM16 (Avisoft Bioacoustics) and transferred to the computer via the UltraSoundGate 116 USB audio device (Avisoft Bioacoustics) using Avisoft RECORDER software (Avisoft Bioacoustics). Following USV measurements, the body temperature of the pups was immediately measured using a surface temperature sensor and reader (Testo) placed on the pup stomach for 30 seconds. Hereafter, pup length and weight were measured. The pup was then placed back in a corner of the home cage to measure maternal retrieval time. In between pups, the glass container was thoroughly cleaned with 70% ethanol and fresh bedding was applied. At P3, after pup USV recording, a small amount of green Ketchum animal tattoo ink (Ketchum Manufacturing inc.) was injected in the paws of each pup for identification purposes, and a small piece of tail was taken for genotyping.

Recorded sound files were analyzed using the SASlab Pro software (Avisoft Bioacoustics). First, a fast Fourier transform was conducted (512 FFT length, 100% frame, Hamming window, and 75% time window overlap) to create spectrograms with 488 Hz of frequency and 0.512 ms of time resolution. Next, section labels were generated using the ‘Automatic Parameter Measurements setup’. Here, the spectral entropy setting was used with a max entropy of 0.4 and a hold time of 30 ms. Identified elements were transferred into labels and each element was manually checked for overlap with the USV, after which the total number of USV was calculated. Peak frequency, duration, amplitude and frequency modulation were then measured at the mean of the element. The amplitude was measured in decibels relative to full-scale (dBFS) where 0 dBFS corresponds to the maximum level of the wave that can be recorded without clipping at the selected gain. All measured sound that does not surpass this is negative relative to this level. The absolute of the measured value in dBFS was then subtracted from 100.

A substantially large number of calls were collected from the ubiquitous and conditional knockout pups over development. The values of all calls were averaged per pup and age, and in all cases the numbers shown in Supplementary Table 1 were used as sample size for the statistical analysis. For the comparison of peak frequency between genotypes, median values were used instead of averages, as the distribution is not Gaussian. Histograms of the average relative abundance of peak frequency per genotype, including 95% confidence intervals (dotted lines) were generated using Graphpad Prism. Due to the variable number of animals used per experiment, a mixed-effects ANOVA with Geisser-Greenhouse correction and Tukey post-hoc test was used unless otherwise noted. Kernel density estimation (KDE) plots were generated with Seaborn v0.12.2 (62) and matplotlib (63). KDE values were scaled based on the number of data points in each dataset, and a common color scale was determined using the minimum and maximum KDE values per feature. Using Plotly (64), interactive 3D scatter plots were generated for each age and genotype. Sequential correlation analysis of the duration of calls with previous calls (N-1) was performed using excel. Peak frequency, duration, amplitude and frequency modulation values were standardized and t-SNE was applied to the scaled data with a random state of 42 and a perplexity of 30. Scaled data was used for clustering via a spectral clustering approach using nearest neighbor affinity with a random state of 42. The characteristic parameters of each call cluster, along with the relative fractions of clusters throughout development or between genotypes were calculated using excel.

### Analysis of single cell RNA dataset

The dataset used in the present study was obtained from the Gene Expression Omnibus (GEO) dataset website (https://www.ncbi.nlm.nih.gov/gds). The dataset (65) was interrogated using the SeuratV3 R package (66). Mitochondrial genes were removed using the ‘PercentageFeatureSet’ tool, and data was transformed using SCTransform. Data was normalized and scaled according to standardized procedures, and variable features (n = 3000, vst) were identified. To identify differential gene markers between ‘Pcdh10 positive’ and ‘Pcdh10 negative’ groups, novel metadata was added to all cells based on their *Pcdh10* count (> 0). By using the ‘FindAllMarkers’ tool of Seurat, differential genes with a fold change of at least 1.5 between both groups were identified. Gene Ontology (GO) over-representation analysis was performed with the ClusterProfiler package (67,68). Gene symbols of identified hub genes and all dataset genes were converted to Entrez IDs using the MyGene package (69), and the Genome wide annotation for Mouse (“org.Mm.eg.db”) database from BioConductor was used as background for the ‘enrichGO’ tool of ClusterProfiler. DotPlots were constructed via the ggplot2 package (70).

### Statistical Analysis

Statistical analysis was performed using GraphPad Prism 9 software (GraphPad Software). Appropriate statistical tests were used depending on the data distribution and experimental design. For normally distributed data, we employed t-tests or ANOVA followed by post-hoc tests for multiple comparisons. Non-parametric tests such as the Mann-Whitney U test or Kruskal-Wallis test were used for non-normally distributed data. In all cases, a p-value less than 0.05 was considered statistically significant. Data are presented as mean ± standard deviation (SD) unless otherwise noted.

## Results

According to our hypothesis, *Pcdh10* might be involved in interneuron development and maturation. To get a better overview of *Pcdh10* expression at earlier developmental stages, *Pcdh10* mRNA was investigated on wild-type (Wt) embryonic (E15.5) coronal paraffin sections using *in situ hybridization* (ISH) (Figure 1A). At this stage, *Pcdh10* can predominantly be found in the subpallium and the ventricular zone (VZ) of the pallium. Within the subpallium, *Pcdh10* was expressed in the VZ and mantle zone (MZ) of the ganglionic eminences (GEs), but clear signal was missing from the subventricular zone (SVZ). Moreover, a salt-and-pepper expression pattern is present, with regions of increased expression in the telencephalon including the striatal region, the insular cortex, the bed nucleus of the stria terminalis, the piriform cortex, the lateral preoptic area (POA) and the amygdala, in particular in the nucleus of the lateral olfactory tract (nLOT). The migratory stream from the pallium towards the amygdala (the ventropallial migratory stream (VMS)) outlines a region with strong *Pcdh10* expression within the subpallium. In the thalamus, the habenula and dorsal thalamic nuclei are clearly marked, in addition to the zona incerta (ZI) and the thalamic reticular nucleus (TRN).

**Figure 1:**
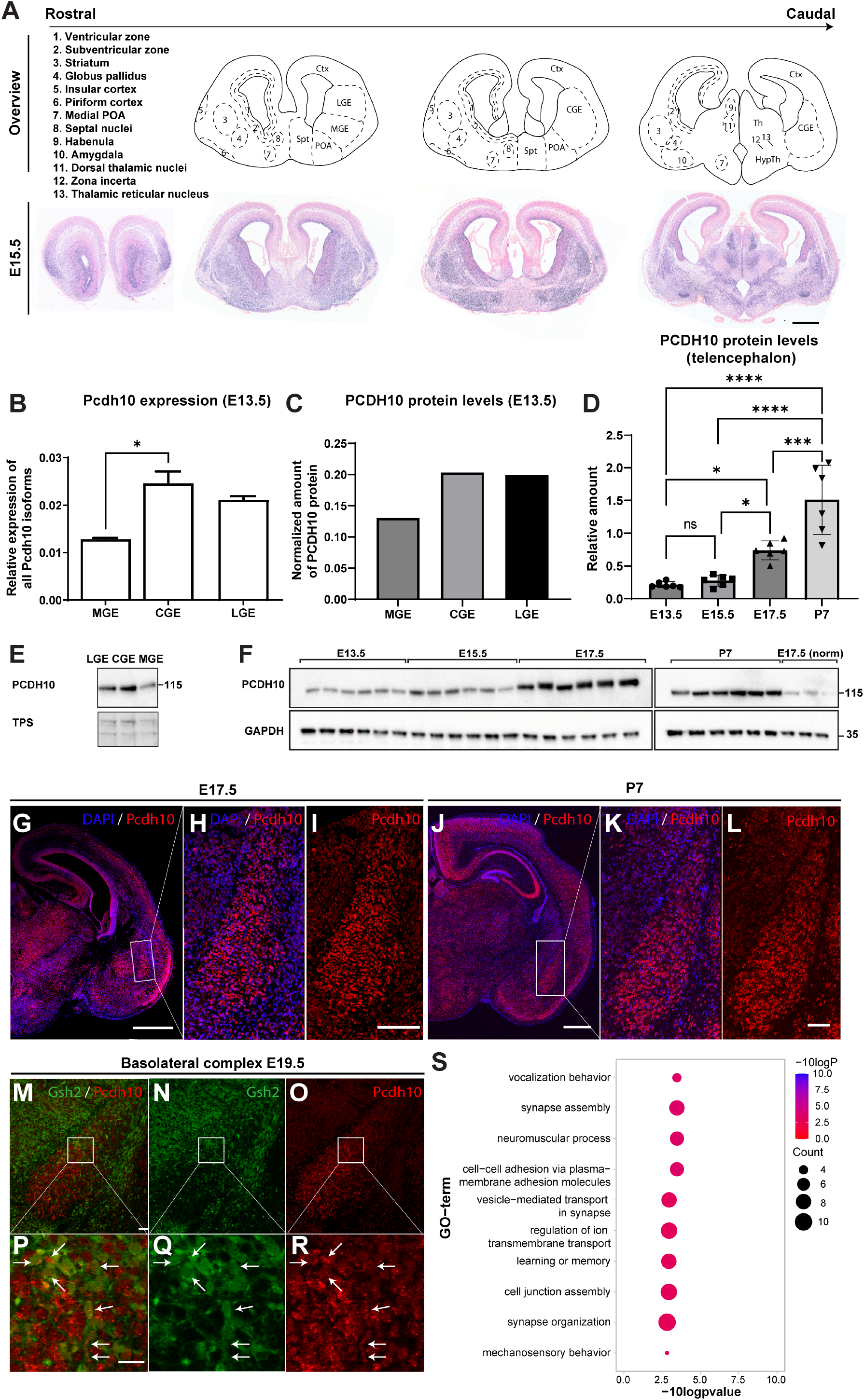
Pcdh10 in the embryonic and juvenile murine wild-type brain. (A) Overview of In Situ Hybridisation against murine Pcdh10 on coronal sections at E15.5. Scalebar 500µm. (B) Relative expression of Pcdh10 in the ganglionic eminences at E13.5. Pcdh10 levels are highest in CGE and LGE. * p ≤ 0.05 as assessed with one-way ANOVA. n=2. (C) Relative protein levels of PCDH10 in pooled samples of the ganglionic eminences at E13.5.Samples are a mix of three biological replicates. n=1. (D) Relative protein levels of PCDH10 in the telencephalon at different stages of development. Proteins levels increase over embryonic and postnatal development. * p ≤ 0.05, *** p ≤ 0.001, **** p ≤ 0.0001, as assessed with ne-way ANOVA. n=6. (E) Blot showing the protein levels of PCDH10 and a total protein stain in pooled samples of the ganglionic eminences at E13.5. TPS = total protein stain. (F) Blot showing the protein levels of PCDH10 and GAPDH in the telencephalon at different stages of development. (G, J) Overview of coronal paraffin brain sections (E17.5 (G), P7 (J)) stained with Hybridisation Chain Reaction against murine Pcdh10.. Scalebar 500µm. (H-I, K-L) Zoom in on basolateral complex (BLC) on coronal brain section (E17.5 (H-I), P7 (K-L)). Scalebar 100µm. (M-O) Overview of the wild-type BLC on coronalsections stained with HCR probes against Pcdh10 (E19.5). Pcdh10 is present in both excitatory and inhibitory neurons. Scalebar 100µm. (P-R) Zoom in on M-O. Scalebar 20µm. (S) Dotplot visualizing the top 10 over-represented Gene Ontology (GO) terms for the ‘biological process’ group for genes identified for the ‘Pcdh10 positive’ and ‘Pcdh10 negative’ groups in adult murine brain (P56). Ctx = Cortex, LGE = Lateral ganglionic eminence, MGE = medial ganglionic eminence, CGE = Caudal ganglionic eminence, POA = Pre-optic area, Spt = Septum, Th = Thalamus, HypTh = Hypothalamus.

The ganglionic eminences (GEs) generate the majority of interneurons in the telencephalon, a cell type that is implicated in ASD etiology. Progenitor cells from the GEs give rise to various subtypes of interneurons, each programmed to mature into interneurons with unique functions and destinations. To investigate whether *Pcdh10* is preferentially expressed in the different GEs, medial (MGE), lateral (LGE) and caudal (CGE) ganglionic eminence tissue was dissected at E13.5 and the mRNA content analyzed. As shown in Figure 1B, normalized expression of *Pcdh10* was highest in the CGE, closely followed by LGE and lowest in MGE. At E13.5, protein levels of PCDH10 in the GEs follow a similar pattern to mRNA expression (Figure 1C,E). To study the dynamics of PCDH10 protein levels during embryonic brain development, and potentially link *Pcdh10* to developmental processes ongoing at different ages, whole telencephalon was collected at E13.5, E15.5, E17.5 and P7 and submitted to Western Blot analysis (Figure 1D,G). PCDH10 levels were found to rise, first steadily at mid-to-late gestational stages (E13.5-E17.5), then sharply between late embryonic stages (E17.5) and early postnatal life (P7), hinting towards a strong involvement in of *Pcdh10* in postnatal development (p = 0,979 (E13.5 – E15.5), p = 0.018 (E13.5 – E17.5), p < 0.0001 (E13.5 – P7), p = 0.042 (E15.5 – E17.5), p < 0.0001 (E15.5 – P7), p = 0.0006 (E17.5 – P7), n = 6). The original western blot images are shown in Supplementary Figure 2.

Previous studies show strong *Pcdh10* expression in the juvenile and adult amygdaloid complex. Our ISH results at E15.5 imply that this expression already emerges during embryonic development. We therefore further investigated this expression at late embryonic and early postnatal developmental stages with HCR (Figure 1G-L). At both E17.5 and P7, the basolateral complex (BLC) can be delineated within the basal forebrain based on the strong signal of *Pcdh10.* At P7, *Pcdh10* is strongly expressed in the basolateral amygdala, with a weak, patchy expression pattern visible in the lateral amygdala (Figure 1J-L). This difference in expression levels is already noticeable at embryonic stages (E17.5) but less pronounced (Figure 1G-I). The BLC of the amygdala is primarily referred to as a pallial- or cortical-like area within the ventral forebrain, with the majority of neurons being excitatory neurons derived from the pallium, and a lower percentage of subpallium-derived inhibitory interneurons (basal amygdala: 22%, lateral amygdala: 16%) (71). To investigate the expression of *Pcdh10* in *Gsh2*-lineage interneurons of the BLC, an HCR was performed at E19.5 in brains in which the *Gsh2*-lineage was labeled with eGFP. *Pcdh10* was expressed both in *Gsh2-*lineage interneurons (white arrows), as well as in other, likely excitatory, cells of the BLC (Figure 1M-R).

To gain an unbiased view on the cell-type specificity and the potential role of cells expressing *Pcdh10* in the mouse brain, a published scRNAseq adult mouse telencephalon dataset was analysed (65). *Pcdh10* was moderately expressed in the cells, with 27.5% of all cells containing at least one count of *Pcdh10*. A method based on virtual differential gene expression analysis via the Seurat function ‘FindAllMarkers’ was then performed. Here, we identified genes that could be considered markers for the group of cells that show at least one *Pcdh10* count versus the group of cells that do not show any counts for *Pcdh10* (cut-off > 0). 49 differentially expressed genes with a fold change of at least 1.5 were identified using this method (Supplementary Table 2). As shown in Figure 1S, highly relevant significant GO-terms of biological process (BP) were identified. The GO term “vocalization behavior” is associated with ASD and links to the observed phenotype of *Pcdh10* mutant mice (34,36). This implies that *Pcdh10* is expressed in cell types involved in control of speech or, alternatively, can be found in a common regulatory network shared with other vocalization-related genes (*Foxp2, Cntnap2, Nrxn1, Nrxn3*). Among the identified markers several genes that are linked to ASD are present, including two neurexins (*Nrxn1* and *Nrxn3*), the receptor-type tyrosine-protein phosphatase T gene (*Ptprt*) and the vocalization gene contactin-associated protein 2 (*Cntnap2*). In addition to vocalization and ASD, multiple over-represented GO terms congregate at the synaptic level.

To investigate whether interneurons are involved in the phenotypes previously observed in *Pcdh10* mutant mice, we specifically knocked out *Pcdh10* in interneurons, by crossing Pcdh10^fl/+^ mice (50) with Gsh2-iCre transgenic mice (52) (Figure 2A). *Gsh2* emerges early during embryonic development (E9.5), selectively labels LGE- and part of CGE-derived interneurons in the telencephalon, and delivers cells eventually populating many of the regions with high *Pcdh10* expression, including the amygdala, striatum and olfactory bulbs (72–75). By crossing this line with a transgenic floxed RCE line (51), all *Gsh2-*expressing cells and their progeny will be green fluorescent. The successful knockout of *Pcdh10* in *Gsh2*-lineage cells was validated in sorted, targeted cells (eGFP^+^) via bulk sequencing at E15.5, and a strong reduction of *Pcdh10* counts could be observed in the telencephalon of cKO embryos as compared to control embryos (Control: 161.60 ± 34.58, cKO: 28.39 ± 14.12, p < 0.0001, n = 6, Figure 2B).

**Figure 2:**
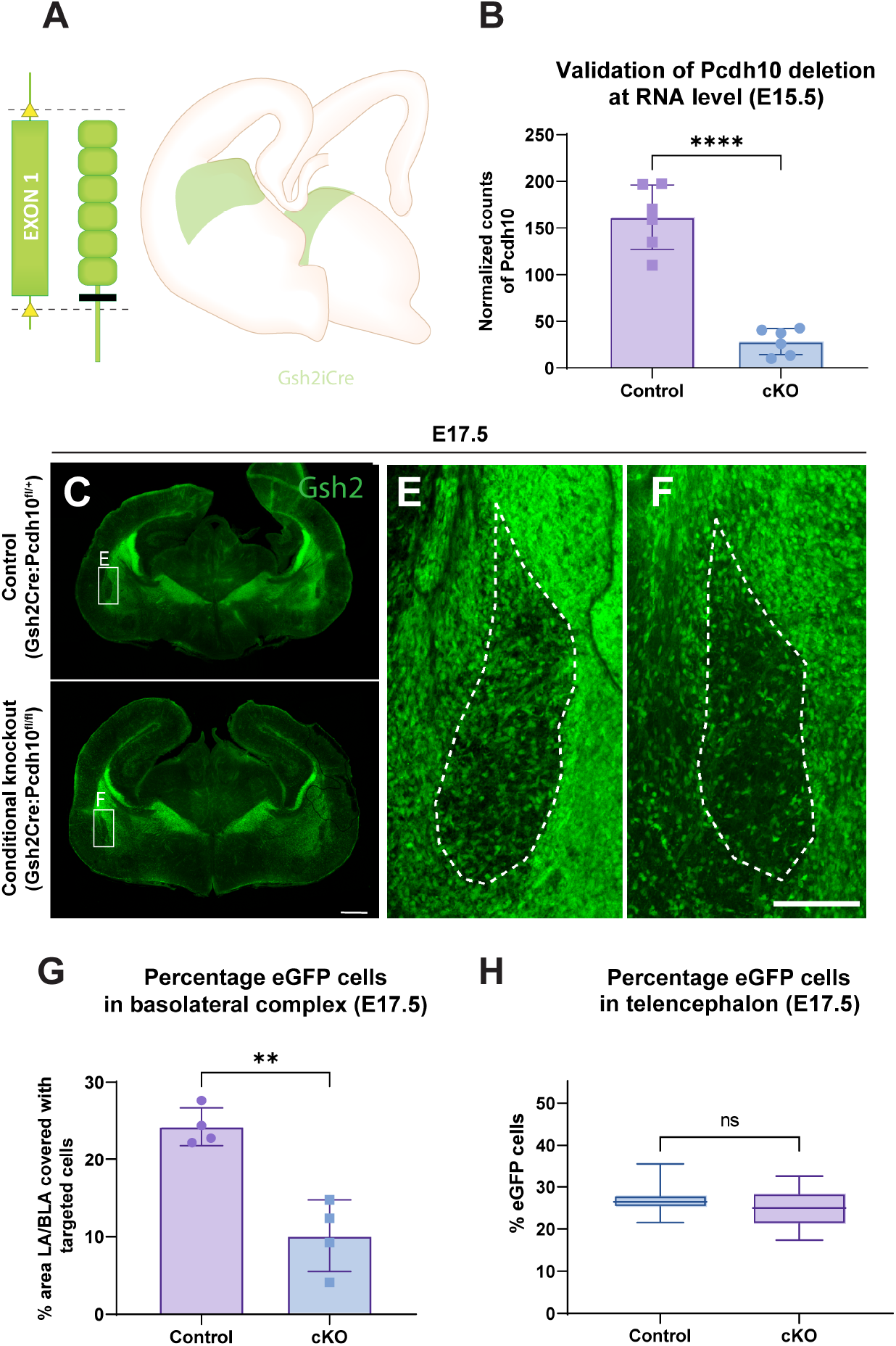
Reduced amount of Gsh2-lineage interneurons in the basolateral complex(E17.5) in a Pcdh10 conditional knockout model. (A) Visual representation of the Pcdh10 conditional knockout model. The first floxed exon of Pcdh10 is excised in targeted cells after crossing with a Cre-driver line. An additional crossing is done with RCE mice to induce green fluorescence in targeted cells. (B) Validation of the Pcdh10 conditional knockout model via bulk RNA sequencing of control and conditional knockout telencephalon at E15.5. Counts are significantly reduced in conditional knockout mice as compared to control mice. (C-D) Overview of Gsh2 expression in coronal vibratome sections (60µm) of control and conditional knockout telencephalon at a caudal level (E17.5). Scalebar 500µm. (E-F) Zoom in on the basolateral complex. Targeted (eGFP^+^) interneurons are reduced in the basolateral complex of the conditional knockout as compared to the control condition. Scalebar 100µm. (G) Quantification of the percentage of basolateral complex area that is occupied by targeted Gsh2-lineage interneurons in control and conditional knockout embryos (E17.5). A significant difference is present between control and conditional knockout, as assessed with two tailed unpaired t-test. ** p ≤ 0.01. n = 4. (H) Percentage targeted (eGFP^+^) cells in the telencephalon of control and conditional knockout embryos at E17.5. No significant differences could be observed as assessed with two-tailed unpaired t-test, p > 0.05. n = 11 (Control) and n = 6 (cKO).

Interneuron dysfunction can contribute to the development of ASD, and *Pcdh10* shows an exceptionally strong expression in the subpallium. A crucial step in elucidating the role of *Pcdh10* in brain development and ASD was to investigate the positioning and amount of interneurons in the Gsh2:cKO model. No broad abnormal positioning of the interneurons could be observed within this model. However, specifically within the BLC, the area occupied by targeted *Gsh2*-lineage interneurons was strongly reduced in cKO brain as compared to control brains (Control: 24.27% ± 2.44, cKO: 10.12% ± 4.61, p = 0.002, n = 4, Figure 2C-G). In addition, the targeted interneurons surrounding the BLC also seemed reduced based on the amount of eGFP signal. To quantify this, we analysed the amount of targeted (*Gsh2*-lineage, eGFP^+^) cells in the entire telencephalon of Gsh2:Control and Gsh2:cKO embryos with flow cytometry at E17.5 (Figure 2H). No significant difference could be observed in the amount of targeted cells between cKO and Control, indicating that this reduced signal in the BLC is not a general result of overall reduced *Gsh2-*lineage interneurons in the telencephalon (p = 0.492).

Previous studies indicated that mice haploinsufficient for *Pcdh10* displayed alterations in socio-affective communication through isolation-induced USV (34,36), a change typically associated with increased anxiety in the pup (76,77). These studies however did not investigate which cells might be responsible for this phenotype, and did not distinguish between potential cell type-specific effects of *Pcdh10* haploinsufficiency. A knockout model and the Gsh2:cKO model were therefore compared to investigate whether altered *Pcdh10* expression in interneurons affects ultrasonic calling, as a measurement for anxiety. To mitigate the effect of the LacZ cassette in the mouse line used by Schoch et al. (2017) (36), a new ubiquitous Pcdh10 knockout (KO) line was generated. To this end, male Pcdh10^fl/+^ mice were crossed with a general maternal Cre line (Pgk1-Cre) to target all cells of the developing progeny (53). After backcrossing for at least three generations, protein was isolated and the loss of PCDH10 was validated in whole body of heterozygous (He) and homozygous (Ho) embryos (E17.5). Heterozygous embryos contained halved PCDH10 levels and homozygous embryos completely lost PCDH10 within the telencephalon (Ho: 0.089 ± 0.031, He: 0.963 ± 0.444, Wt: 2.328 ± 0.935, p = 0.025 (Wt – He), p = 0.001 (Wt – Ho), p = 0.152 (He – Ho), one-way ANOVA, n = 4, Figure 3A,B). The original western blot is again shown in Supplementary Figure 2.

**Figure 3:**
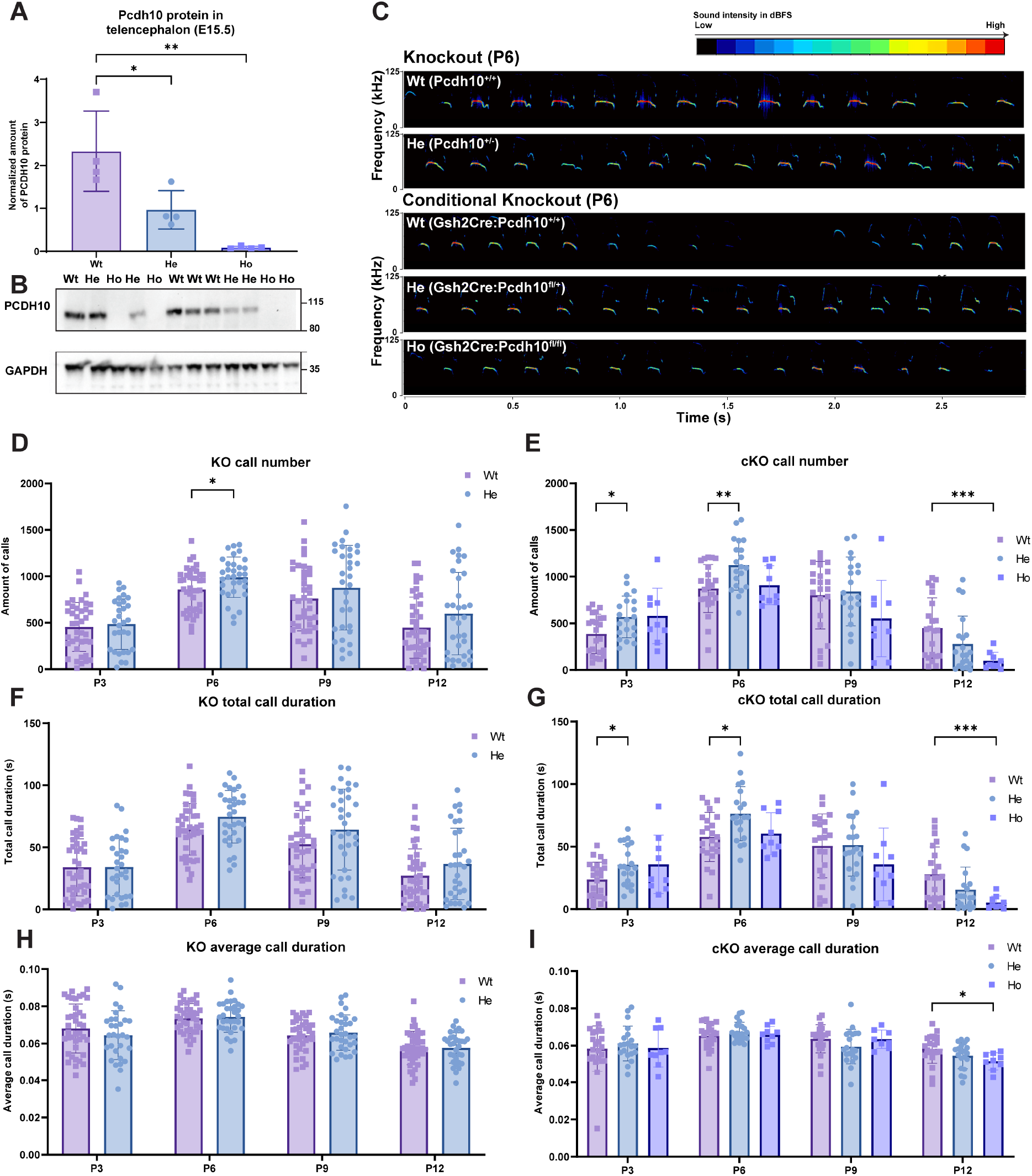
Abnormal ultrasonic vocalizations (USVs) in heterozygous and homozygous Pcdh10 knockout pups between P3-P12. (A-B) Validation of Pcdh10 knockout line via Western Blot (P7). The amount of protein is significantly reduced in heterozygous and homozygous pups as assessed with one-way ANOVA. n = 4. (C) Representative spectrograms of USV of a P6 wild-type, heterozygous Pcdh10KO, and heterozygous and homozygous conditional knockout littermates. (D - E) Amount of isolation-induced USV emitted by wild-type and heterozygous Pcdh10KO pups (E) and by wild-type, heterozygous and homozygous Gsh2-lineage Pcdh10 conditional knockout pups (F). USV of heterozygous KO are increased at P6 and USV of heterozygous cKO pups are increased at P3 and P6 as compared to wild-type pups. USV of homozygous cKO pups are decreased at P12 as compared to wild-type pups. (F - G) Total call duration of isolation-induced USV in wild-type and heterozygous Pcdh10KO pups (F) and in wild-type, heterozygous and homozygous Gsh2-lineage Pcdh10 conditional knockout pups (G). Total call duration of heterozygous Pcdh10KO pups is increased at P6 and total call duration of heterozygous cKO pups are increased at P3 and P6 as compared to wild-type pups. Total call duration of the USV of homozygous cKO pups are decreased at P12 as compared to wild-type pups. (H - I) Call duration of isolation-induced USV in wild-type and heterozygous Pcdh10KO pups (H) and in wild-type, heterozygous and homozygous Gsh2-lineage Pcdh10 conditional knockout pups (I). Call duration of heterozygous Pcdh10KO pups is similar to wild-type pups and call duration of homozygous cKO pups is decreased at P12 as compared to wild-type pups. * p ≤ 0.05, ** p ≤ 0.01, *** p ≤ 0.001.

Pups of both lines were monitored for 15 days following birth and all pups showed normal weight gain, growth and body temperatures (Supplementary Figure 3). Development but not genotype contributed to changes in these parameters (aging p < 0.0001). Isolation-induced USV were measured every third day from P3-P12 and a representative spectrogram of a P6 Pcdh10^+/+^ (KO Wt), Pcdh10^+/-^ (KO He), Gsh2Cre^+/+^:Pcdh10^X/X^ (Gsh2:cKO Wt), Gsh2Cre^+/-^:Pcdh10^+/fl^ (Gsh2:cKO He) and Gsh2Cre^+/-^:Pcdh10^fl/fl^ (Gsh2:cKO Ho) pup is shown in Figure 3C. Across test days, more than 300000 isolation-induced USV were recorded and analyzed. Total call number was assessed for all lines and genotypes (Figure 3D,E). In the *Pcdh10* KO line, genotype affected USV numbers (p = 0.037). Specifically, pups haploinsufficient for *Pcdh10* (KO) showed increased USV emission rates as compared to their wild-type littermates, significant at P6 when these calls peak and persisting until P12 (P6: p = 0.041, Figure 3D). No developmental shift in the period of USV production could be identified (p age*genotype > 0.05). In contrast, the developmental trajectories of USV numbers differed between genotypes for the cKO line (p age*genotype < 0.0001). Similar to haploinsufficient KO pups, heterozygous cKO pups showed increased levels of isolation-induced USV as compared to wild-type littermates, observable as early as P3 and peaking at P6 (P3: p = 0.025; P6 p = 0.010; Figure 3E). At these ages, the difference between heterozygous cKO pups and wild-type littermates even surpassed those observed between heterozygous *Pcdh10* KO pups and controls. By P9, this difference became less pronounced, leveling off to wild-type levels by P12. Notably, call numbers from homozygous and heterozygous cKO pups were comparable at P3, but values from homozygous pups thereafter decreased, resulting in comparable numbers to wild-type pups at P6 and decreased numbers at P9 and P12, respectively (P12: p = 0.0002 (Wt – Ho)). Total call duration revealed similar trends for the ubiquitous and conditional knockout mice (Figure 3F,G). In general, call duration changed throughout development but was unaffected by genotype, although the developmental trajectory of the duration of calls differed between genotypes of the cKO (age p < 0.0001, cKO age*genotype p = 0.0001, Figure 3H,I). In addition to the reduced number of calls between wild-type and homozygous cKO pups at P12, call duration was decreased (p = 0.038 (Wt - Ho), Figure 3I).

Previous studies revealed that differences in USV emission rates and acoustic call features, including amplitude, peak frequency and frequency modulation, can affect maternal retrieval behavior (13). Due to the increased specificity of the experimental manipulation and the more prominent effect of *Pcdh10* deletion in *Gsh2-*lineage interneurons on isolation-induced USV, we opted to further investigate acoustic call features in the conditional knockout line. Peak frequency, call duration and frequency modulation all changed throughout development (age p-value for peak frequency p = 0.0008, call duration p < 0.0001, frequency modulation p < 0.0001). At P6, calls were emitted at lower median peak frequencies in heterozygous as compared to wild-type pups (p = 0.032, Figure 4A), while the frequency modulation was increased (p = 0.028 (Wt – He), Figure 4B). The amplitude of calls was unchanged between genotypes (Figure 4C). Moreover, sequential analysis of the call duration revealed that there is a positive correlation between each call (N) and the previous call (N – 1) (Figure 4D). This correlation was relatively high at P3 and P6 (r > 0.4) and then decreased as development progressed, suggesting a high level of stereotypic call emission at early developmental stages. The developmental progression of these correlation values are affected by genotype in the conditional knockout (p age*genotype = 0.041). Heterozygous cKO pups exhibit a steeper decrease of correlation values over development as compared to wild-type littermates from P9 onward, potentially affecting their ability to communicate with their mother or littermates (P9 p = 0.040 (Wt – He)).

**Figure 4:**
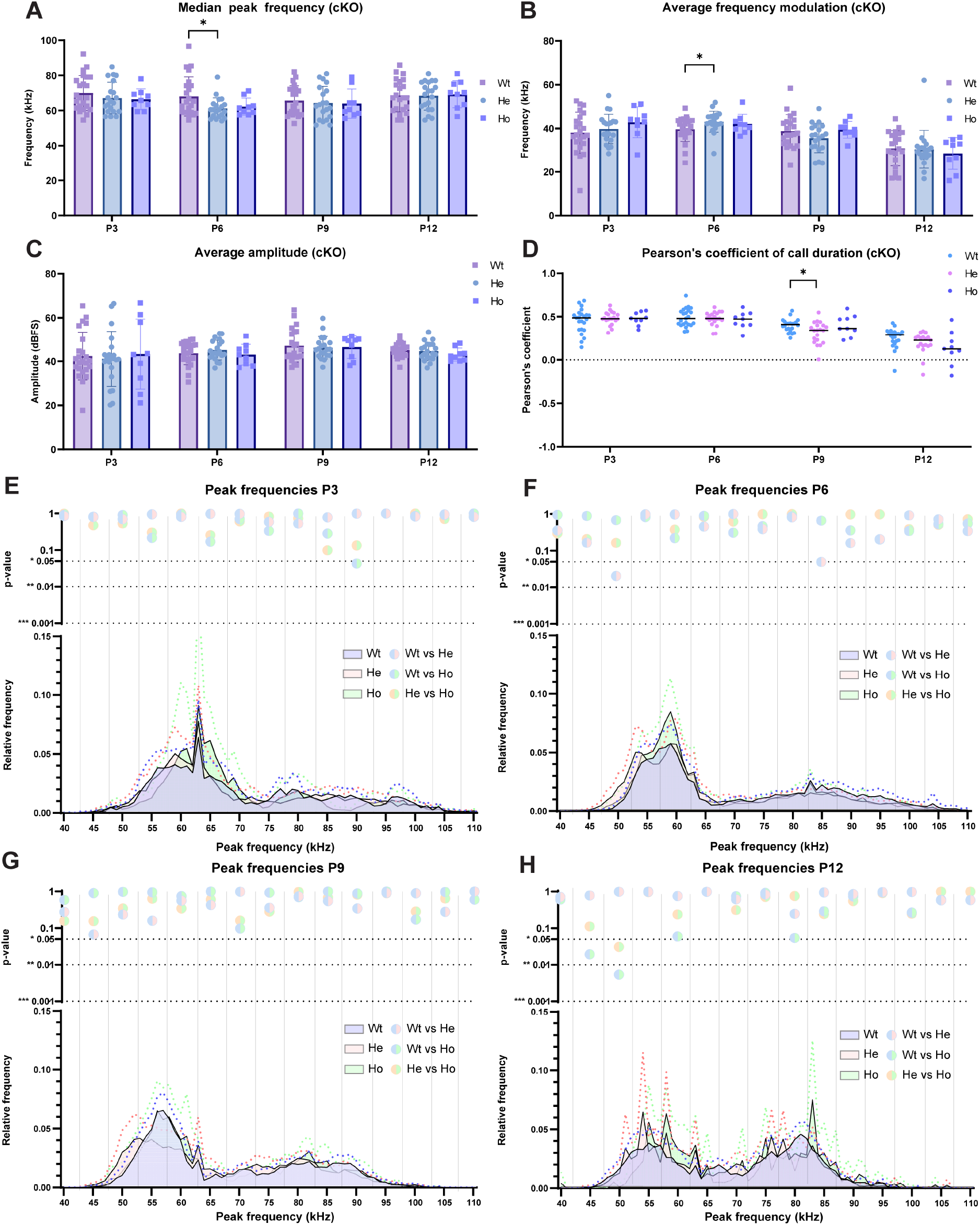
Pcdh10 conditional knockout pups exhibit shift in peak frequency and amplitude in ultrasonic vocalizations (USV). (A) peak frequency, (B) frequency modulation and (C) amplitude of isolation-induced USV in wild-type, heterozygous and homozygous Gsh2-lineage Pcdh10 conditional knockout pups. Peak frequency of the USV of heterozygous pups is decreased at P6 as compared to wild-type pups, while frequency modulation is increased. USV amplitude is unchanged between genotypes. (D) Pearson’s coefficient of call duration (N-1) of isolation-induced USV in wild-type, heterozygous and homozygous Gsh2-lineage Pcdh10 conditional knockout pups. Correlation reduces throughout development. Pearson’s coefficient is significantly reduced in heterozygous pups as compared to wild-type littermates at P9. (E – H) Histograms showing the relative fraction versus peak frequency of isolation-induced USV in wild-type, heterozygous and homozygous Gsh2-lineage Pcdh10 conditional knockout. P-values of the grouped relative frequency bins are shown. * p ≤ 0.05, ** p ≤ 0.01, *** p ≤ 0.001.

Different call types have previously been identified in isolated rat and mouse pups. To classify the different types of isolation-induced USV, we further analysed the more than 130000 isolation-induced calls emitted by the wild-type, heterozygous and homozygous conditional knockout pups over their development. Histograms with binned p-values of the relative fraction of peak frequency values averaged by pup were created and analysed for each age and genotype (Figure 4E-H). This allowed us to observe differences that were not evident from purely analyzing averaged or median values, as peak frequency is not normally distributed. As shown, two main bands of peak frequency can be observed throughout development, including one band around 60 kHz (55 – 65 kHz) and one broader band around 80 kHz (70 – 100 kHz), from now on referred to as low pitch and high pitch USV. Over development, the peak frequency of both bands decreased and the spread of values in both bands became less prominent. In addition, the relative contribution of the high pitch calls (∼ 80 kHz) increased with age. When comparing wild-type, heterozygous and homozygous conditional knockout pups, subtle differences in peak frequencies can be observed. The relative contribution of the low pitch calls is slightly increased for the homozygous cKO pups at earlier stages (P3-P6), but normalizes after the first postnatal week. Visible from P6 onward is the slight decrease of the peak frequency of calls emitted by heterozygous cKO pups as compared to wild-type and homozygous littermates, significantly different for the 47.5 kHz – 52.5 kHz bin (p = 0.020) and 82.5 kHz – 87.5 kHz bin (p = 0.049). This difference is still visible at P9, but not significant, and has disappeared by P12.

To better understand the distribution of calls with certain peak frequency, duration, amplitude and frequency modulation values over development and between genotypes, kernel density plots and 3D interactive plots were generated (Figure 5 and Interactive Supplementary Files). In Figure 5A-L, wild-type profiles of peak frequency versus the other parameters are shown from P3-P12. Clearly visible in all plots is the previously described emergence of high pitch calls as the pups age, evolving from a diffuse cluster to a more distinct cluster (top half of Figures 5A-L). In general, the range of the duration of calls decreased as development progressed (P3: 0.01s – 0.11s; P12: 0.03s – 0.10s, Figure 5A-D). Interestingly, the frequency modulation of calls at early postnatal stages (P3) is distributed across a large range (0 - 80 kHz) but evolves towards two major groups of frequency modulation at later stages (P6 onward, 0 - 30 kHz and 40 - 65 kHz, Figure 5E-H). Similar to the duration, the variation in amplitude across all calls becomes less extreme over time, with the majority of calls situating between 20 dBFS and 80 dBFS at P3 and between 30 dBFS and 70 dBFS at P12 (Figure 5I-L).

**Figure 5:**
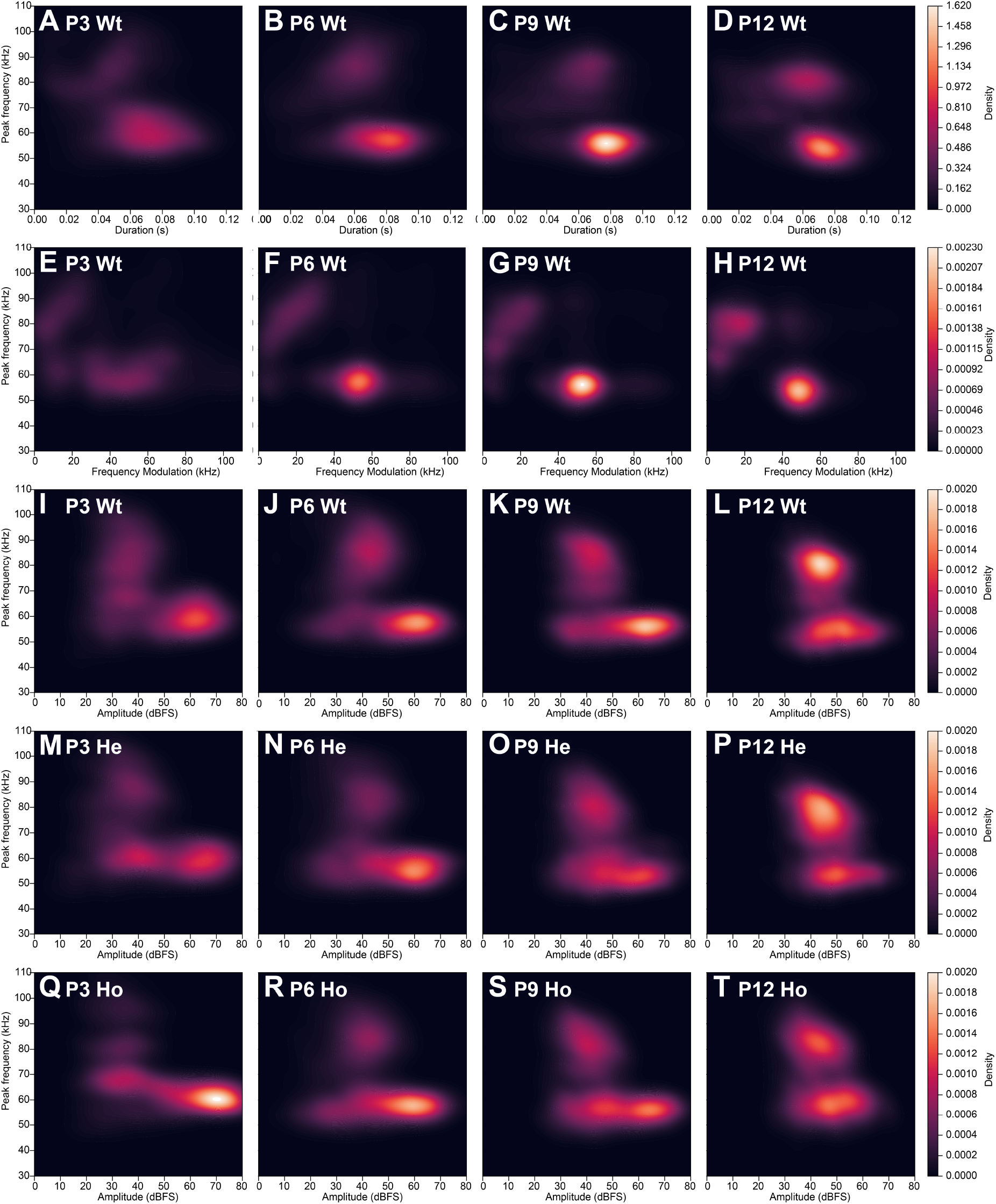
Call clusters change throughout development and between conditional knockout genotypes. (A-L) Kernel density estimation plots of peak frequency (kHz) versus duration (s, A-E), frequency modulation (kHz, E – H) and amplitude (dBFS, I-L) of wild-type pups throughout development (P3, P6, P9, P12). Pattern and density of call clusters change throughout development. (I-T) comparison of kernel density estimation plots with peak frequency (kHz) versus amplitude (dBFS) showing identified call clusters and densities of Gsh2-lineage conditional knockout wild-type (Wt, I - L), heterozygous (He, M - P) and Homozygous (Ho, Q - T) pups at postnatal days (P)3 (I, M, Q), P6 (J, N, R), P9 (K, O, S) and P12 (L, P, T).

The two call types defined by their peak frequency each show their own preferential duration, amplitude and frequency modulation. In general, low pitch calls show a slightly increased duration and are louder as compared to high pitch calls (Figure 5A-D, I-L). Moreover, low pitch calls show a medium-to-high frequency modulation, evident from P6 onward when frequency modulation becomes more refined (40 - 65 kHz) (Figure 5E-H). In contrast, the majority of high pitch calls have low frequency modulation values at all ages (0 - 30 kHz).

When comparing between genotypes, call amplitudes show a shift towards louder calls in the homozygous cKO pups at P3 (Figure 5I, M, Q). Likewise, we observe a subtle relative increase of calls at lower peak frequency, together with a relatively decreased amount at the higher frequency band in the homozygous condition, with the heterozygous condition showing an intermediate state (Supplementary Figure 4). No overt differences in the call duration were observed in kde plots of all genotypes (Supplementary Figure 4A-L). Notably, in the homozygous condition at P3 high-pitch calls show further separation based on peak frequency, with a band around 80 kHz and a band around 100 kHz instead of the more continuous peak frequency range in the heterozygous and wild-type condition (Supplementary Figure 4M, 4Q, 4U, Interactive Supplementary Files).

To better illustrate the existence of different call types, calls were clustered into six clusters via a spectral clustering algorithm. The t-SNE plot and the average cluster values for peak frequency, call duration, amplitude and frequency modulation are shown in Figure 6A-E. Within the low pitch calls, three clusters can be further identified (cluster 0, 4 and 5). Calls belonging to cluster 5 are the most rare at all ages and are characterized by extremely low frequency modulation (6.2 kHz ± 1.7), short duration (0.034s ± 0.005) and low amplitudes (32.48 dBFS ± 2.52), illustrating the emission of low-pitched, soft, short and simple sounds (Interactive Supplementary Files, Figure 6B-E). The other two subclusters (0 and 4) within the lower pitch group show the typical medium-to-high frequency modulation (52.7 kHz ± 4.7 and 50.2 kHz ± 2.6, respectively), but vary in duration and amplitude, both showing relatively high values (duration: 0.085s ± 0.005 and 0.068s ± 0.004; amplitude: 62.75 dBFS ± 2.93 and 45.47 dBFS ± 2.42). These clusters illustrate the existence of more complex, lower-pitched sounds. The higher frequency band can be subdivided into two main clusters (2 and 3) that show similar peak frequencies at all ages (see Interactive Supplementary Files, Figure 6B). Cluster 3 is the loudest within the high-pitch call group (43.80 dBFS ± 4.42) with medium duration (0.060s ± 0.003) but relatively low frequency modulation (18.5 kHz ± 2.4). A minority of calls within this cluster extend to lower frequency ranges. Instead, calls of cluster 2 are soft, simple calls with a low amplitude (39.85 dBFS ± 2.63), extremely short call durations (0.025s ± 0.003) and very low frequency modulation values (8.2 kHz ± 1.5). One cluster (1) spans the entire frequency range and is characterized by lower amplitudes (32.48 dBFS ± 3.86), medium duration (0.066s ± 0.005) but extremely high frequency modulation (66.6 kHz ± 5.0), hinting at the emission of relatively quiet but intense and complex sounds. Throughout development, all clusters remain present although the relative contribution of each the cluster differs throughout development (age*cluster p < 0.0001, Figure 6F, Supplementary Figure 5). Moreover, peak frequency, call amplitude and frequency modulation change in a cluster-dependent manner throughout development (Cluster*age p value peak frequency < 0.0001, duration p = 0.011, amplitude p < 0.0001, frequency modulation p < 0.0001). As higher-pitched calls become more prevalent over time, the extension of cluster 3 and 1 into lower frequency ranges diminishes (Interactive Supplementary files). This increase of high pitched-calls can be mainly attributed to an increase of calls belonging to cluster type 2 and 3 and a decrease of calls belonging to clusters 0 and 1 (Figure 6B-E).

**Figure 6:**
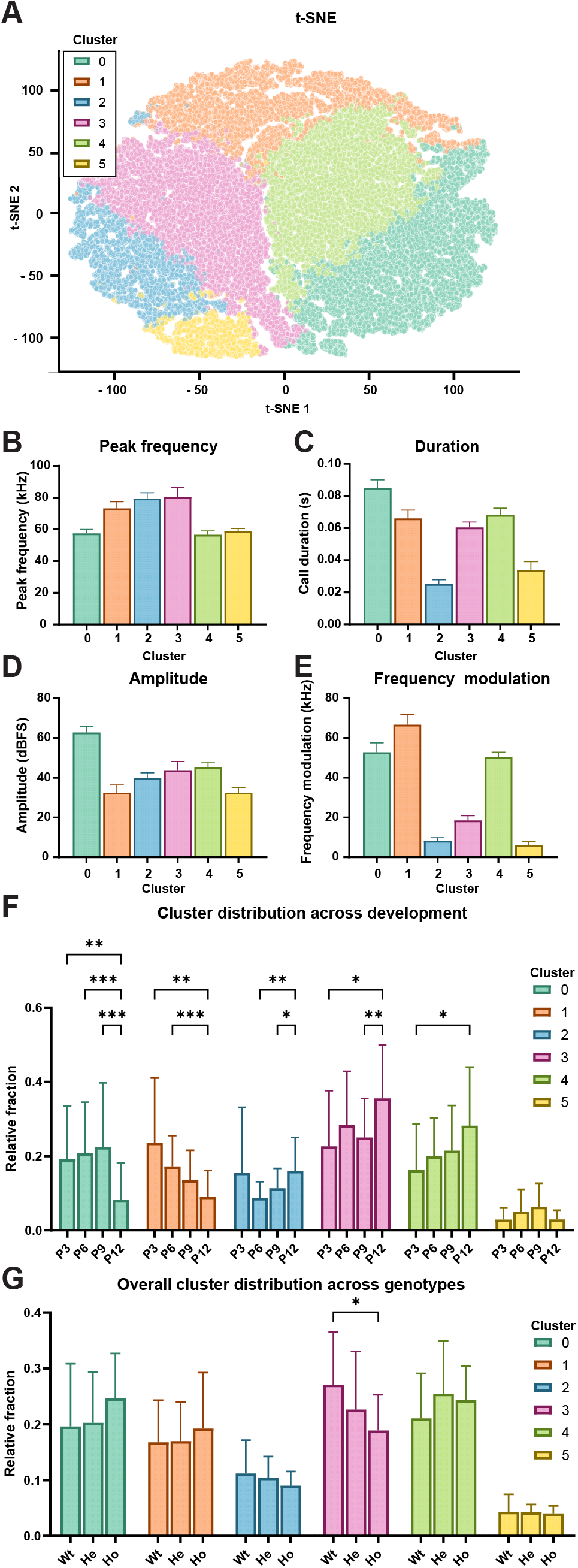
Dynamic appearance of different call types throughout development and impact of conditional Pcdh10 loss. (A) t-SNE plot of all calls labeled by their respective cluster. (B-E) Average peak frequency (B), duration (C), amplitude (D) and frequency modulation (E) values per cluster. (F) The relative fraction of wild-type calls belonging to a cluster over development (P3-P6-P9-P12). (G) The relative fraction of calls from wild-type, heterozygous and homozygous conditional knockout pups per cluster at all ages. * p ≤ 0.05, ** p ≤ 0.01, *** p ≤ 0.001.

When comparing the distribution of the call clusters between genotypes over development, some slight overall differences can be observed (Figure 6G, p cluster 3 = 0.0257 (Wt – Ho)). Nonetheless, no genotype effects were significant when analyzing the ages individually (Supplementary Figure 6). To investigate whether loss of *Pcdh10* affected socio-affective communication with the mother, maternal retrieval time was analysed in our conditional knockout model. In this model, the mother is always wild-type. As some pups tended to independently crawl back to their nest, especially at later developmental stages, retrieval latency was only taken into account if they were retrieved by their mother. Developmental age and genotype directly affected the retrieval time, with an additional shift in the retrieval time developmental trajectory by genotype (p age > 0.0001, p genotype = 0.013, p age*genotype = 0.0002, Supplementary Figure 7).

## Discussion

Here, we investigated the expression and function of *Protocadherin10* (*Pcdh10*) in the developing rodent brain, a cell adhesion molecule linked to the development of ASD in humans (78). We utilized a combination of interneuron-specific conditional and full *Pcdh10* knockout mice to zoom in on certain neuronal cell populations, anxiety and socio-affective communication. In conditional knockout model, we observed reduced interneurons in the basolateral complex of the amygdala and altered socio-affective communication, implicating interneurons in ASD etiology.

### Throughout development Pcdh10 is expressed in brain regions linked to ASD and social behavior

Involvement of members of the non-clustered Protocadherin family in proliferation, sorting and spatial patterning of the brain has become evident over recent years (79–82). Non-clustered protocadherins are associated with neural circuit formation (83), and *Pcdh10* has been directly implicated in wiring of olfactory and amygdalar circuitry (34,36,84,85). The combinatorial expression of cadherin superfamily members, including protocadherins, delineates amygdala subdivisions, and this was suggested to underlie the functional organization of the amygdala (86). We found *Pcdh10* to be preferably expressed in developing structures of the basal ganglia, olfactory and limbic systems. Brain regions that show high *Pcdh10* expression overlap significantly with regions functionally linked to emotional regulation, fear and social behavior (87–93). Perturbed expression of *Pcdh10* could therefore result in circuit-level dysfunctions manifesting as ASD-associated behavior.

The presence of *Pcdh10* in these brain structures already at early development stages suggests its role is not confined to maintenance of neural circuits but also extends to the early establishment of these structures. PCDH10 levels are already high at E13.5 and continue to rise during embryonic and early postnatal development, pointing towards an increasingly important role during neurogenesis, neuronal migration, axonal outgrowth and synapse formation and elimination. Expression of *Pcdh10* within synapses of the amygdala at juvenile and adult stages was previously reported (34,36), and *Pcdh10* was linked to synapse elimination in hippocampal cell cultures (94). Our data shows that *Pcdh10* is already strongly expressed in the BLC of the amygdala during embryonic development (E17.5), but that the preferential expression within the BLA becomes stronger after birth. This dynamic change could imply that *Pcdh10* is involved in early development of the entire BLC, but is then restricted to the BLA to regulate circuit formation and maintenance. Abnormal synapse formation or maintenance have been implicated in ASD pathophysiology (78,95). Synaptic dysfunction and improper E/I balance both affect neuronal transmission, suggesting that disrupted communication between key neuronal groups is fundamental to ASD development. At juvenile stages, both excitatory and inhibitory synapses in the BLC express *Pcdh10* (34).

### Reduced amount of interneurons in the basolateral complex of the Gsh2:cKO mouse model

Our study confirmed the expression of *Pcdh10* predominantly in LGE and CGE at embryonic stages. Strong expression was also observed in some structures located in the ventral telencephalon, including the BLC of the amygdala. The amygdala consists of several nuclei that could be considered predominantly “pallial” or “subpallial” in origin (96). Previous studies have focused on the role of *Pcdh10* in excitatory neurons of the BLC during postnatal stages. However, our results indicate that already at embryonic stages, both excitatory neurons and G*sh2*-lineage interneurons express *Pcdh10* in the BLC.

Loss of *Pcdh10* within this subpopulation of interneurons caused a significant reduction in their numbers in the BLC of the amygdala at E17.5. In this conditional knockout model, the excitatory cells of the BLC remain positive for *Pcdh10* while *Gsh2*-lineage interneurons lose this expression. This targeted loss specifically in interneurons might have compromised integration into the *Pcdh10-*positive BLC. Previous studies have revealed a strong association between several non-clustered protocadherins and sorting defects observed both *in vitro* and in the developing brain, including the formation of specific aggregates due to differential δ2-protocadherin expression *in vitro* (81), sorting anomalies and reduced interneuron migration in the cortex of *Pcdh19* mosaic females (81,83) and the mislocalization of *Gad1-*expressing cells in the thalamus of *Pcdh10b* knockdown zebrafish (97). These findings collectively suggest that loss of protocadherin expression in adjacent cells might affect their positioning and circuit integration. Alternatively, a proliferative or apoptosis defect in the progenitor/mature interneuron population could also explain the reduced interneuron numbers. In cancer, *Pcdh10* has been linked to both these processes (25), but in the telencephalon we could not observe an overall reduction of *Gsh2-*lineage interneurons at E17.5.

### Ultrasonic vocalizations

A reduced number of interneurons could potentially disturb the balance between excitation and inhibition and impair BLC functioning, which we measured using USV as a readout. Isolation-induced USV are sometimes compared to the crying of human babies, as both serve a similar purpose (98). In infant individuals diagnosed with ASD and in mouse models, abnormal crying and isolation-induced USV have been observed (11,12,98,99). While reduced pup ultrasonic calling is typically interpreted as evidence for socio-affective communication impairment (8,11,12), increased emission rates are often interpreted as an indirect measure of enhanced anxiety levels (76,77).

An inverted U-shaped developmental call emission pattern could be identified for both the KO and the cKO, as previously described in mice (16–18). Hoshina et al. (2022) has reported decreased USV emission in homozygous Pcdh10 KO as compared to wild-type pups (34). They postulated that the increased USV previously observed by Schoch et al. (2017) in the heterozygous knockout mice might be caused by non-specific effects of the LacZ cassette (34,36). However, in our study both full body as well as conditional heterozygous models also exhibited heightened USV emission rates, most prominent during the peak call emission period (P3 – P6). In contrast, homozygous Gsh2:cKO pups showed decreased instead of increased pup ultrasonic callings at P12, revealing a dynamic and more complex involvement of *Pcdh10*. Future studies should investigate whether ubiquitous heterozygous and homozygous knockout pups show varying USV phenotypes. A majority of the genes linked to ASD are described as gene-dosage sensitive genes, and point mutations and/or copy number variations (including both deletions and duplications) affecting only one allele are common in individuals with ASD (78). For some genes, both individuals with deletions and those with duplications have been described to suffer from ASD, although the specific symptoms can be distinct (78) and transcriptomic changes related to ASD might appear at different life stages (100). These findings highlight that the function of ASD-related genes might be dynamic throughout life and might be gene-dosage sensitive.

Previous studies on isolation-induced USV in mice and rats showed that calls can be divided into distinct clusters of call subtypes based on their peak frequency, call duration, amplitude and frequency modulation (11,17,101– 103). Mothers were observed to show categorical frequency perception for certain frequency bands, indicating that emission of calls with abnormal peak frequency might affect care and retrieval behavior (13). We identified 6 call subtypes (clusters), defined by peak frequency, duration, amplitude and frequency modulation, which could provide additional insight into the infant mouse communicative potential.

Calls cluster in two peak frequency ranges, including low pitch calls around 60 kHz (55 – 65 kHz) and high pitch calls around 80 kHz (70 – 100 kHz). Low pitch calls were predominantly longer, more complex and louder as compared to high pitch calls. In previous experiments in mice and rats, the high pitch call type was described to have low frequency modulation and call duration, similar to our call clusters 2 and 3, while the two low pitch call types could be divided into one with low frequency modulation and one with high frequency modulation, similar to our cluster 0 and 4 versus 5 (102,104). We have further subdivided these call types and identified an additional group with high pitch and high frequency modulation (cluster 1) in CD-1 mice. The prevalence of the high pitch calls increased over development, as also previously observed (16,17,102,103,105). These two peak frequency clusters are also clearly present in pups heterozygous for *Pcdh10*KO and *Pcdh10*cKO, although the peak frequencies are subtly shifted towards lower values with increased frequency modulation in pups heterozygous for conditional knockout of *Pcdh10* in *Gsh2-*lineage interneurons. Clusters become more comparable to wild-type patterns as development progresses. The behavioral implications for these changes remain largely unknown, but genotype affected the retrieval behavior of the mother. Since the contribution of the more complex isolation-induced calls decreases over development, they might primarily serve to induce a rigid retrieval response in the mother as compared to calls with lower frequency modulation.

In mammals, multiple brain regions are involved in vocal communication, including the anterior cingulate cortex, the periaqueductal gray (PAG), the reticular formation, the motor cortex, basal ganglia, cerebellum and phonatory motoneurons (106). In the conditional *Pcdh10* knockout model, we observed reduced interneurons in the BLC of the amygdala. Changes in the socio-affective communication are more pronounced in the conditional knockout model as compared to the full knockout, implicating the targeted interneuron subpopulation in this phenotype. Abnormal interneuron function and decreased gamma-aminobutyric acid (GABA) transmission in the amygdala directly affects sociability and fear behavior in rodents (47,107,108). In addition, USV have been linked to both anxiety and the amygdala (76,77,109–111). In mice, the amygdala was shown to affect ultrasonic vocalization behavior in the context of courtship via connections to the PAG, which weighs the USV-promoting signals from the preoptic area against the USV-suppressing, GABAergic signals emerging from the amygdala (112). Reduction of GABAergic input from the amygdala, for example due to reduced interneurons as observed in our model, could therefore directly affect the gating of vocalization behavior in the PAG.

Analysis of the adult brain scRNAseq dataset revealed that *Pcdh10*-expressing cells differentially expressed genes involved in vocalization behaviors, including *Foxp2, Cntnap2, Nrxn1 and Nrxn3,* as compared to cells that did not express *Pcdh10.* This might indicate that *Pcdh10* generally functions in brain regions linked to vocalization. *Gsh2* is also strongly expressed in the hindbrain at early developmental stages (113), potentially targeting some of the known vocalization nuclei. Investigation of these regions should reveal whether the observed phenotype primarily is the result of reduced *Pcdh10* in the interneurons of the basal ganglia and amygdala, or if other (hindbrain) regions are involved.

## Limitations

Studies on USV in *Pcdh10* knockout lines have previously been performed, and similar results were obtained in heterozygous pups of the *Pcdh10^LacZ/+^* line used by Schoch et al., (2017) (36). Although the overall statistical significance and difference between the number of calls in heterozygous knockout pups and wild-types is rather modest (at P6 p = 0.041), these findings replicated previously published data and can therefore be considered robust. We did not test ubiquitous homozygous *Pcdh10* knockout pups, complicating comparison with the recent publication by Hoshina et al. (2022) (34). The effects on USV numbers were similar between our tested models, with the heterozygous cKO pups showing an exaggerated effect as compared to the ubiquitous heterozygous KO pups. This is demonstrated by the very low p-values for the age*genotype interaction term (p < 0.0001) and modest p-values for the wild-type versus heterozygous comparison at P6 (p = 0.010). Combined with the robust (replicable) effect seen in the ubiquitous knockout, we can conclude that the conditional knockout shows a similar or stronger effect on USV emission.

The analysis of distinct clusters of call subtypes revealed similar findings as previously published; however we further divided calls into six different subtypes. We want to highlight that this spectral clustering of call types is likely still an underrepresentation of the vocal repertoire of isolated pups, and that some clusters can be further divided into subclusters. The clusters identified in this publication can be used as a general classification of call types based on selected parameters, and serve as a guide for studies that want to further characterize the communicative potential of isolated neonatal pups.

While we show a robust effect of the conditional loss of *Pcdh10* on the amount of interneurons in the amygdala (Control: 24.27% ± 2.44, cKO: 10.12% ± 4.61, p = 0.002), we do not claim or prove that this specific reduction in the amygdala is the major cause of the altered isolation-induced USV behavior in the conditional knockout model. As mentioned in the discussion, *Pcdh10* is co-expressed with other genes involved in vocalization and might be strongly expressed in *Gsh2-*lineage interneurons of other vocalization nuclei. Nonetheless, the specificity of the conditional knockout model proves that inhibitory interneurons are involved in this phenotype, and highlights that *Pcdh10* expression can regulate the ratio of excitatory and inhibitory neurons in at least the BLC. An altered excitation/inhibition balance in the amygdala has previously been linked to altered sociability and anxiety in rodents.

## Conclusions

This study highlights the involvement of *Pcdh10* in anxiety-related behavior and socio-affective communication in developing mice pups and implicates interneuron subpopulations in the behavioral alterations relevant to ASD. *Pcdh10* is strongly expressed in interneurons and expression increases over embryonic and early postnatal development. Targeted deletion of *Pcdh10* in *Gsh2*-lineage interneurons revealed a reduced presence of this subpopulation in the basolateral complex of the amygdala as compared to controls. Isolation-induced USV were altered in heterozygous pups, which exhibited increased emission rates and altered acoustic features including decreased peak frequencies and enhanced frequency modulation. This phenotype is similar and even stronger than that of a ubiquitous heterozygous knockout of *Pcdh10.* Together with the identification of distinct clusters of call subtypes and their subtle variations between genotypes, this emphasizes the complex role of *Pcdh10* in vocal communication and anxiety, two aspects of behavior often affected in ASD.

## List of abbreviations

ASD: Autism Spectrum Disorder
BLA: Basolateral Amygdala
BLC: Basolateral complex
BP: Biological process
CGE: Caudal ganglionic eminence
cKO: Conditional knockout
Ctx: Cortex
dBFS: Decibels relative to full-scale
E: Embryonic day
GABA: Gamma-aminobutyric acid
GEs: Ganglionic eminences
GO: Gene ontology
HCR: Hybridization chain reaction
He: Heterozygous
Ho: Homozygous
HypTh: Hypothalamus
ISH: In situ hybridization
KDE: Kernel density estimation
KO: knockout
LA: Lateral amygdala
LGE: Lateral ganglionic eminence
MGE: Medial ganglionic eminence
MZ: Mantle zone
nLOT: Nucleus of the lateral olfactory tract
NRTC: Non-reverse transcriptase control
NTC: Non-template control
P: Postnatal day
PAG: Periaqueductal grey
PBS: Phosphate-buffered saline
Pcdh10: Protocadherin10
PCR: Polymerase chain reaction
PFA: Paraformaldehyde
POA: Preoptic area
PVDF: Polyvinylidene difluoride
qPCR: quantitative polymerase chain reaction
Spt: Septum
SVZ: Subventricular zone
Th: Thalamus
TRN: Thalamic reticular nucleus
USV: Ultrasonic vocalization
VMS: Ventropallial migratory stream
VZ: Ventricular zone
Wt: Wild-type
ZI: Zona incerta

## Declarations

## Ethics approval

All animal experiments were approved by the KU Leuven ethical committee, in compliance with current Belgian and European regulations (projects P028/2017, P020/2021 and P003/2022).

## Consent for publication

Not applicable.

## Availability of data and materials

All data is available in this published article and the supplementary files. Raw data will be made available upon request.

## Competing interests

The authors declare that they have no competing interests.

## Funding

This project was supported by the Fonds Wetenschappelijk Onderzoek – Vlaanderen (FWO; Research Foundation – Flanders) research and infrastructure projects G0B5916N and I013018N to E.S., G0E6722N and I007022N to M.W. and 1S42720N to LM; by KU Leuven (Internal Funds KU Leuven) through a C14/16/049 project grant and a KA/16/020 small infrastructure grant to E.S. and a BOFZAP Starting Grant to M.W. (PXF-E0120-STG/20/062). Part of the resources and services used in this work were provided by the VSC (Flemish Supercomputer Center), funded by the Research Foundation - Flanders (FWO) and the Flemish Government.

## Authors’ contributions

T.A. performed all experiments with the help of A.B. (amygdala phenotyping), L.G. and A.S. (Western blotting). T.A. analysed all data and L.M. helped with writing scripts for python. A.P. contributed to setting up the mouse colony and critical discussions. A.F. helped with genotyping of all mice. F.V.R provided the floxed *Pcdh10* mouse line and the probe for ISH. T.A. wrote the manuscript. M.W. and E.S. conceptualized the project, contributed feedback for analysis and reviewed and edited the manuscript’s content. All authors contributed to the manuscript and approved the submitted version.

## Acknowledgements

The authors wish to thank Evelien Herinckx for taking care of all laboratory animals used in this study. In addition, we would like to thank Ruth Styfhals, Ali Elagoz and Melissa Chen for help with the weaning of newborn mice. We are grateful to all members of the Seuntjens, Arckens and Moons research groups for critical discussions.

## Additional files

Interactive File 1 _ P3 Wt 3Dscatter.html

Interactive File 2 _ P3 He 3Dscatter.html

Interactive File 3 _ P3 Ho 3Dscatter.html

Interactive File 4 _ P6 Wt 3Dscatter.html

Interactive File 5 _ P6 He 3Dscatter.html

Interactive File 6 _ P6 Ho 3Dscatter.html

Interactive File 7 _ P9 Wt 3Dscatter.html

Interactive File 8 _ P9 He 3Dscatter.html

Interactive File 9 _ P9 Ho 3Dscatter.html

Interactive File 10 _ P12 Wt 3Dscatter.html

Interactive File 11 _ P12 He 3Dscatter.html

Interactive File 12 _ P12 Ho 3Dscatter.html

Interactive files are available at: https://github.com/Tania-Seuntjenslab/Interactive-Files_USV

## Notes

https://github.com/Tania-Seuntjenslab/Interactive-Files_USV

### Competing Interest Statement

The authors have declared no competing interest.

