## Supplemental Data for "Altered Socio-Affective Communication and Amygdala Development in mice with Protocadherin10-deficient Interneurons"

***Supplementary Table 1: Number of pups per age and genotype.***

| | *Strain* | *Age* | | | |
|---|---|---|---|---|---|
| | | **P3** | **P6** | **P9** | **P12** |
| *KO* | Wt (Pcdh10+/+) | 39 | 40 | 39 | 40 |
| | He (Pcdh10+/+) | 30 | 32 | 32 | 32 |
| | Wt (Gsh2iCre+/+;Pcdh10$^{x/x}$;RCE$^{+/-}$) | 23 | 23 | 22 | 22 |
| *cKO* | He (Gsh2iCre$^{+/-}$;Pcdh10$^{fl/+}$;RCE$^{+/-}$) | 20 | 20 | 20 | 20 |
| | Ho (Gsh2iCre$^{+/-}$;Pcdh10$^{fl/fl}$;RCE$^{+/-}$) | 9 | 9 | 9 | 9 |

***Supplementary Table 2: Genes enriched in the Pcdh10-expressing population***

| | **LOG2FC** | **FRACTION IN PCDH10$^{+}$ CELLS** | **FRACTION IN PCDH10$^{-}$ CELLS** |
|---|---|---|---|
| **PCDH10** | 1,993005609 | 1 | 0 |
| **ETL4** | 0,84014181 | 0,651 | 0,379 |
| **SGCZ** | 0,811978626 | 0,663 | 0,4 |
| **GM28928** | 0,807359467 | 0,349 | 0,19 |
| **PDE10A** | 0,768942677 | 0,773 | 0,488 |
| **RBFOX1** | 0,768809346 | 0,82 | 0,562 |
| **CNTN4** | 0,752889419 | 0,698 | 0,403 |
| **MEG3** | 0,746310338 | 0,868 | 0,672 |
| **CDH18** | 0,743389746 | 0,602 | 0,35 |
| **MGAT4C** | 0,741817761 | 0,632 | 0,362 |
| **GRM8** | 0,728555495 | 0,485 | 0,248 |
| **GABRG3** | 0,716311744 | 0,66 | 0,374 |
| **SNHG11** | 0,714501065 | 0,866 | 0,676 |
| **HS3ST4** | 0,698509459 | 0,427 | 0,248 |
| **NRXN3** | 0,695843505 | 0,824 | 0,552 |
| **LHFPL3** | 0,693044637 | 0,555 | 0,308 |
| **KCNQ5** | 0,690478727 | 0,639 | 0,392 |
| **NRG1** | 0,681483835 | 0,778 | 0,544 |
| **ASIC2** | 0,677383816 | 0,784 | 0,5 |
| **FGF14** | 0,672602305 | 0,887 | 0,603 |
| **NXPH1** | 0,658786296 | 0,309 | 0,18 |
| **DPP10** | 0,646562724 | 0,723 | 0,419 |
| **FRMPD4** | 0,640768959 | 0,729 | 0,455 |
| **AGBL4** | 0,638322646 | 0,763 | 0,462 |
| **GALNTL6** | 0,635570554 | 0,573 | 0,359 |
| **SLC35F1** | 0,634040991 | 0,636 | 0,356 |
| **PTPRT** | 0,633940979 | 0,568 | 0,329 |
| **CNTNAP2** | 0,632615113 | 0,797 | 0,526 |
| **CSMD1** | 0,628609434 | 0,858 | 0,579 |
| **GPC6** | 0,617769878 | 0,518 | 0,281 |
| **BRINP3** | 0,615003236 | 0,666 | 0,41 |
| **NTM** | 0,612944548 | 0,829 | 0,512 |
| **FGF12** | 0,612616744 | 0,848 | 0,58 |
| **KCNIP4** | 0,612360185 | 0,808 | 0,584 |
| **GRIP1** | 0,60939857 | 0,67 | 0,403 |
| **PDE7B** | 0,601597222 | 0,465 | 0,287 |
| **PPP2R2B** | 0,600392923 | 0,85 | 0,517 |
| **PTPRN2** | 0,596211046 | 0,784 | 0,484 |
| **RIMS2** | 0,595135032 | 0,827 | 0,54 |
| **PDE4D** | 0,594777482 | 0,845 | 0,605 |
| **GALNT13** | 0,588461382 | 0,672 | 0,397 |
| **FOXP2** | 0,586708002 | 0,345 | 0,179 |
| **SYT1** | 0,586594495 | 0,804 | 0,563 |
| **CACNA2D3** | 0,584972081 | 0,633 | 0,417 |
| **SGCD** | 0,583914291 | 0,554 | 0,328 |
| **NRXN1** | 0,583806268 | 0,926 | 0,684 |

| | LOG2FC | FRACTION IN PCDH10⁺ CELLS | FRACTION IN PCDH10⁻ CELLS |
|---|---|---|---|
| **ADGRB3** | 0,582793364 | 0,915 | 0,667 |
| **ARPP21** | 0,58235211 | 0,7 | 0,459 |
| **CNTN5** | 0,581367845 | 0,623 | 0,397 |

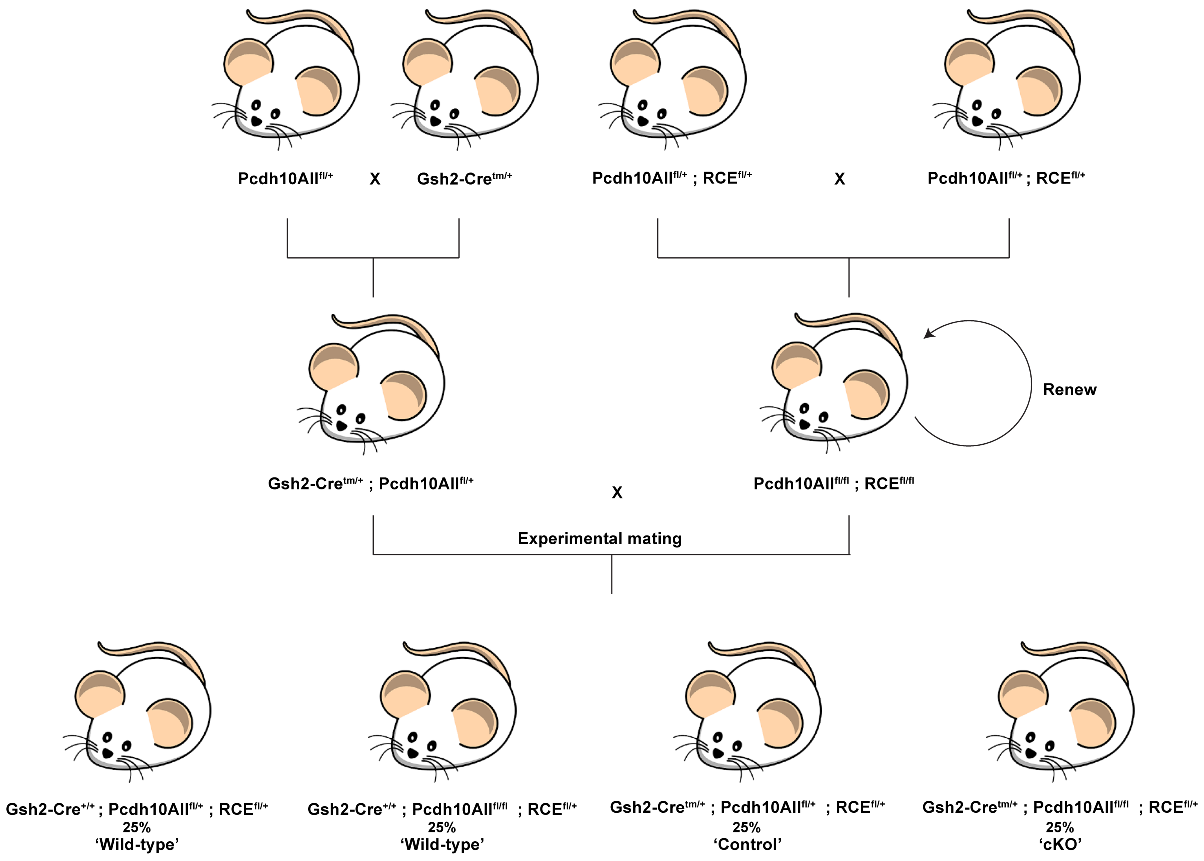


***Supplementary Figure 1: Breeding scheme of*** Pcdh10 ***conditional knockout mice.***

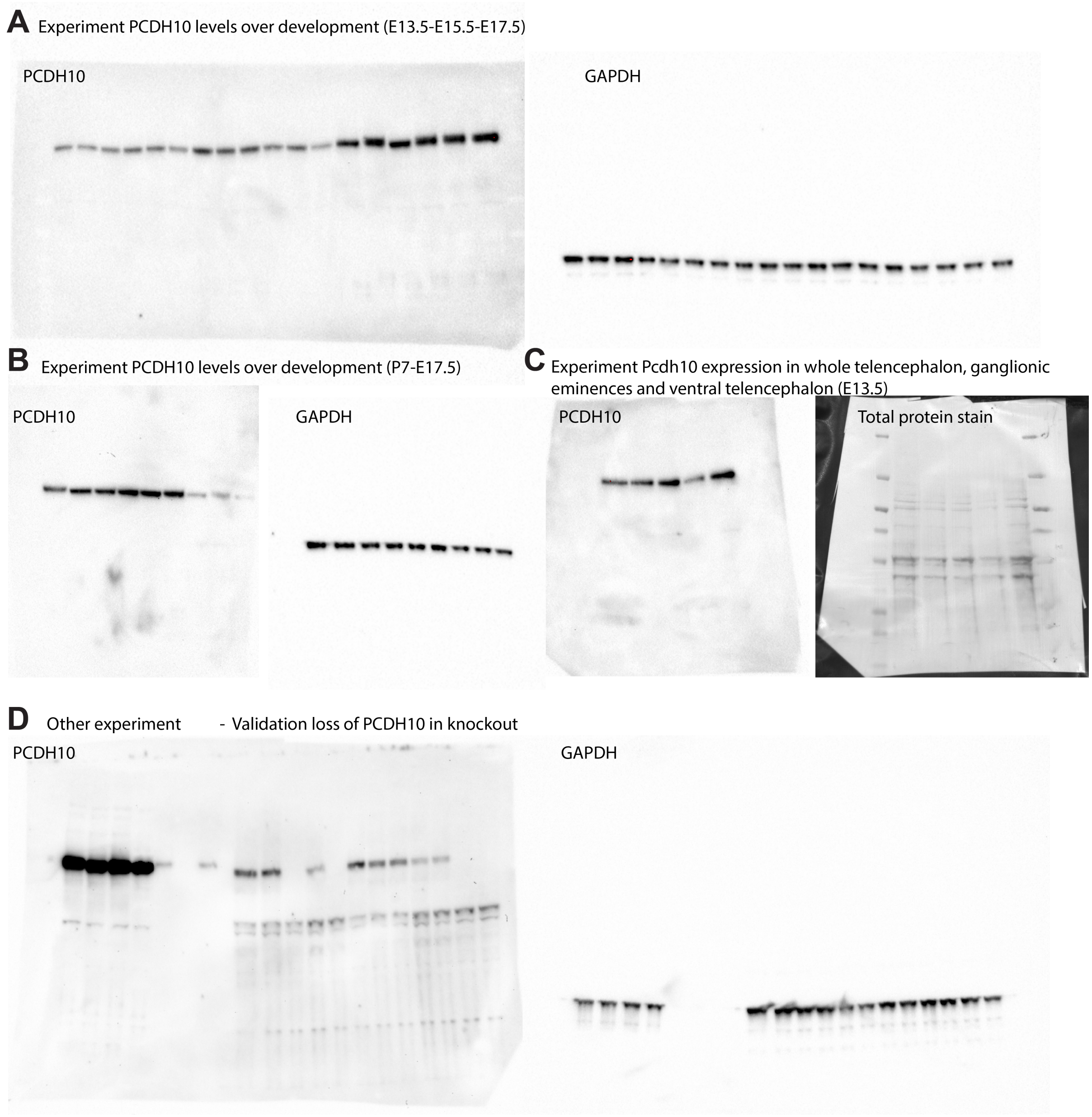


***Supplementary Figure 2: Western blots of experiments in manuscript.***

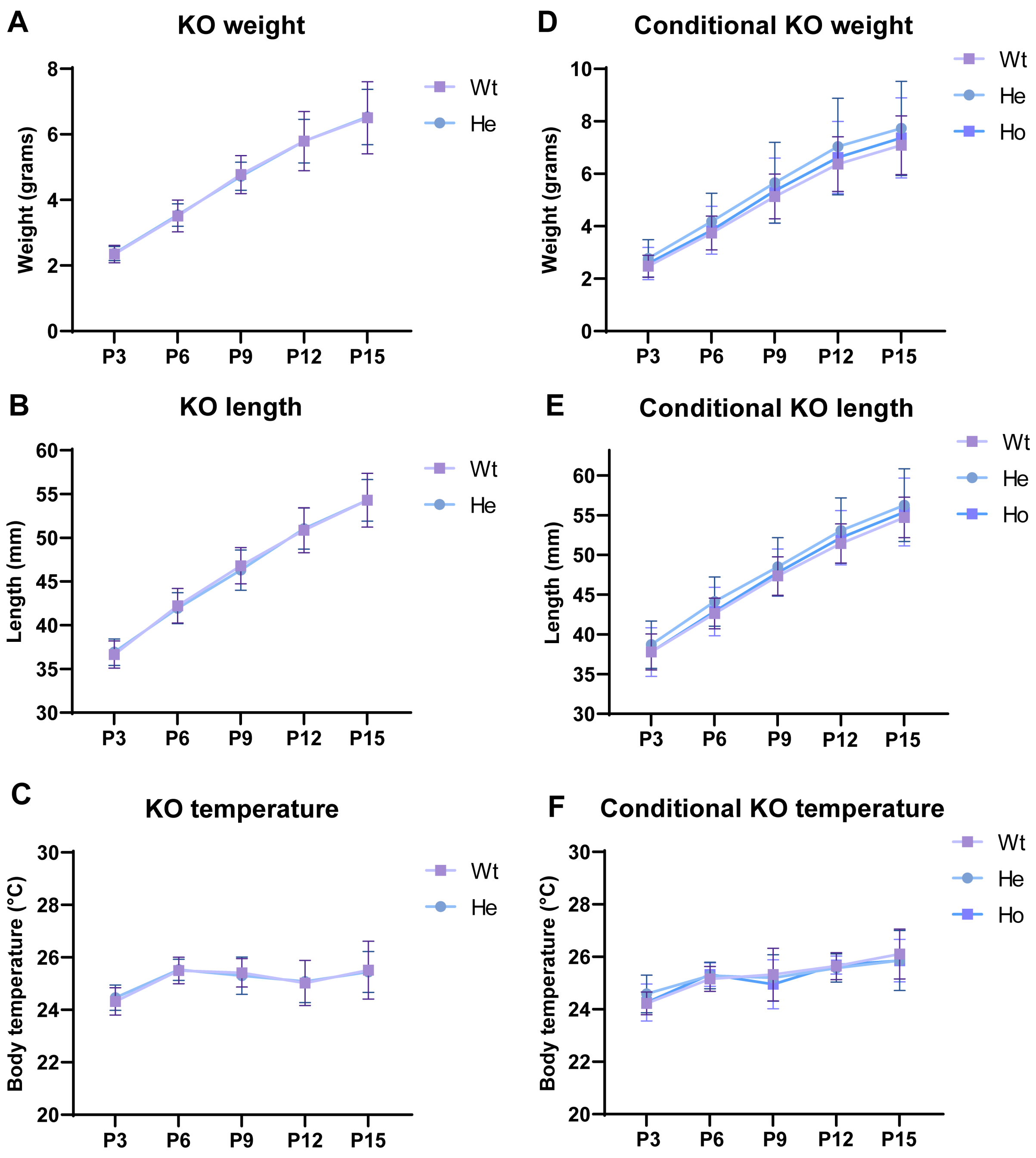


***Supplementary Figure 3: Development of Pcdh10 ubiquitous and conditional knockout pups.*** *Bodyweight (A, D), length (B, E) and temperature (C, F) over early postnatal development (P3-15) for the Pcdh10 ubiquitous (A-C) and conditional knockout (D-F) line.*

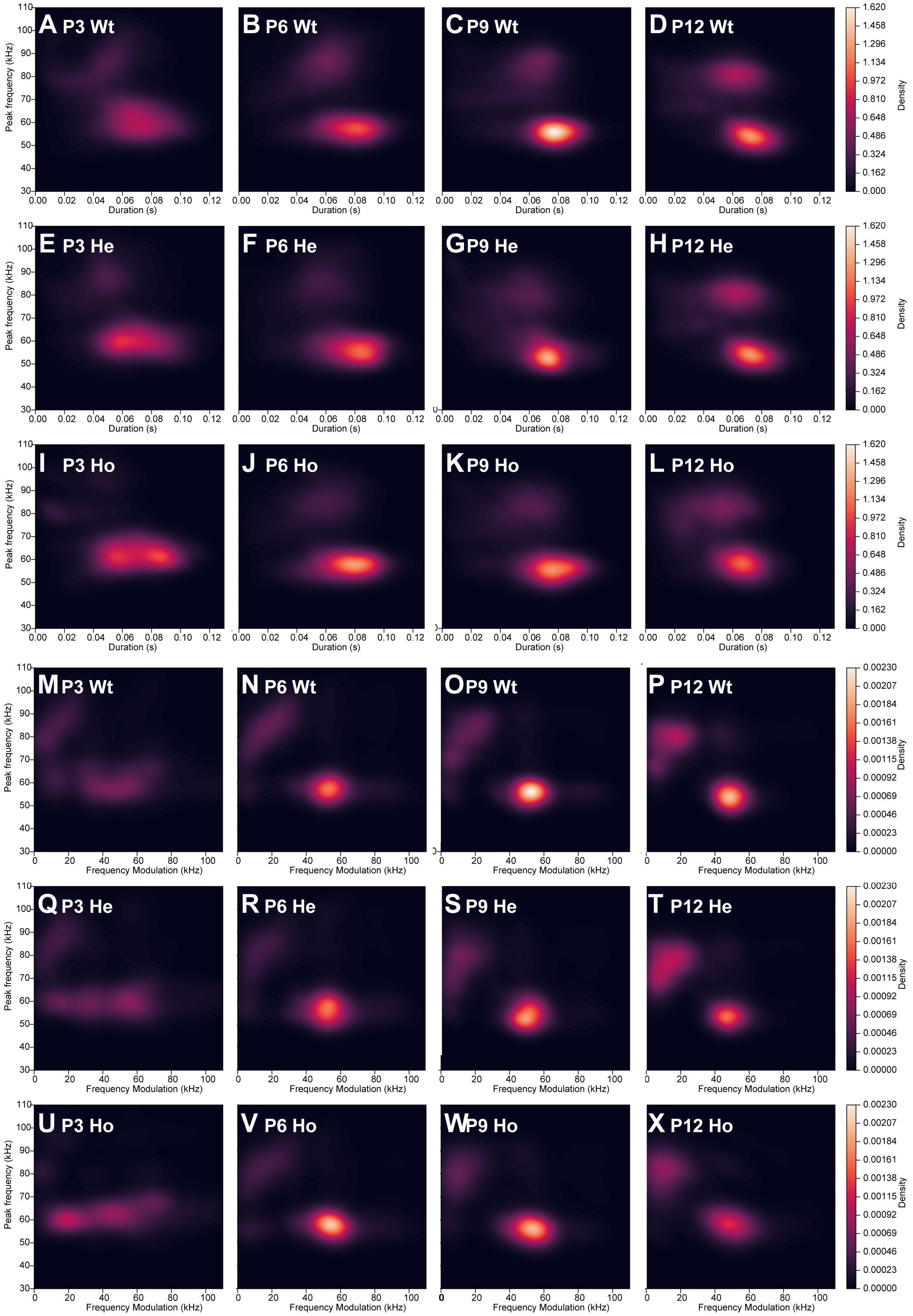


***Supplementary Figure 4: Kernel density estimation plots of call duration and frequency modulation of calls emitted by conditional knockout pups over development (P3-P12).*** *(A-L)Kernel density estimation plots of peak frequency (kHz) versus duration for Wt (A-D), heterozygous (E-H) and homozygous (I-L) pups. (M-X) Kernel*

*density estimation plots of peak frequency (kHz) versus frequency modulation for Wt (M-P), heterozygous (Q-T) and homozygous (U-X) pups.*

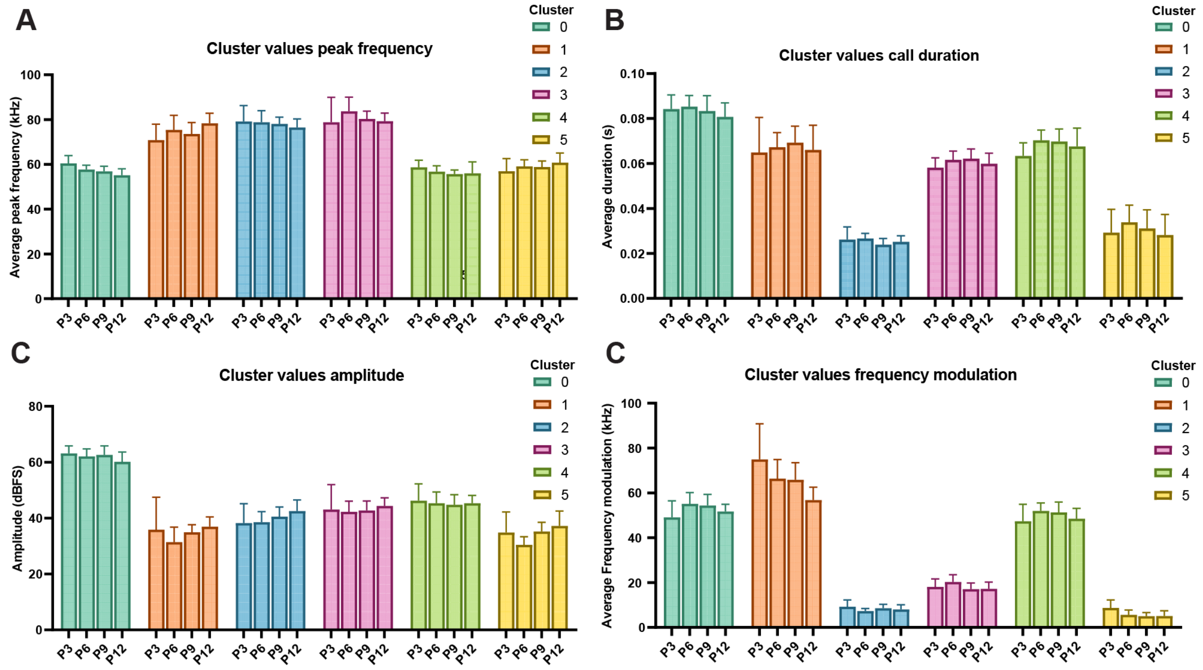


**E**

**p values of different parameters between call clusters**

| p-values | Peak frequency | | | | | | Duration | | | | | |
|---|---|---|---|---|---|---|---|---|---|---|---|---|
| **Cluster** | 0 | 1 | 2 | 3 | 4 | 5 | 0 | 1 | 2 | 3 | 4 | 5 |
| **0** | | | | | | | | | | | | |
| **1** | **** | | | | | | **** | | | | | |
| **2** | **** | **** | | | | | **** | **** | | | | |
| **3** | **** | **** | ns | | | | **** | **** | **** | | | |
| **4** | ns | **** | **** | **** | | | **** | ns | **** | **** | | |
| **5** | ns | **** | **** | **** | * | | **** | **** | **** | **** | **** | |

| p-values | Amplitude | | | | | | Frequency modulation | | | | | |
|---|---|---|---|---|---|---|---|---|---|---|---|---|
| **Cluster** | 0 | 1 | 2 | 3 | 4 | 5 | 0 | 1 | 2 | 3 | 4 | 5 |
| **0** | | | | | | | | | | | | |
| **1** | **** | | | | | | **** | | | | | |
| **2** | **** | **** | | | | | **** | **** | | | | |
| **3** | **** | **** | **** | | | | **** | **** | **** | | | |
| **4** | **** | **** | **** | ns | | | ** | **** | **** | **** | | |
| **5** | **** | ns | **** | **** | **** | | **** | **** | * | **** | **** | |

***Supplementary Figure 5: Values for selected parameters over early postnatal development (P3-P12).*** *Average peak frequency (A), duration (B), amplitude (C) and frequency modulation (D) values of conditional knockout pups (all genotypes) per cluster over early postnatal development (P3-P12). (E) p-values of multiple comparisons.*

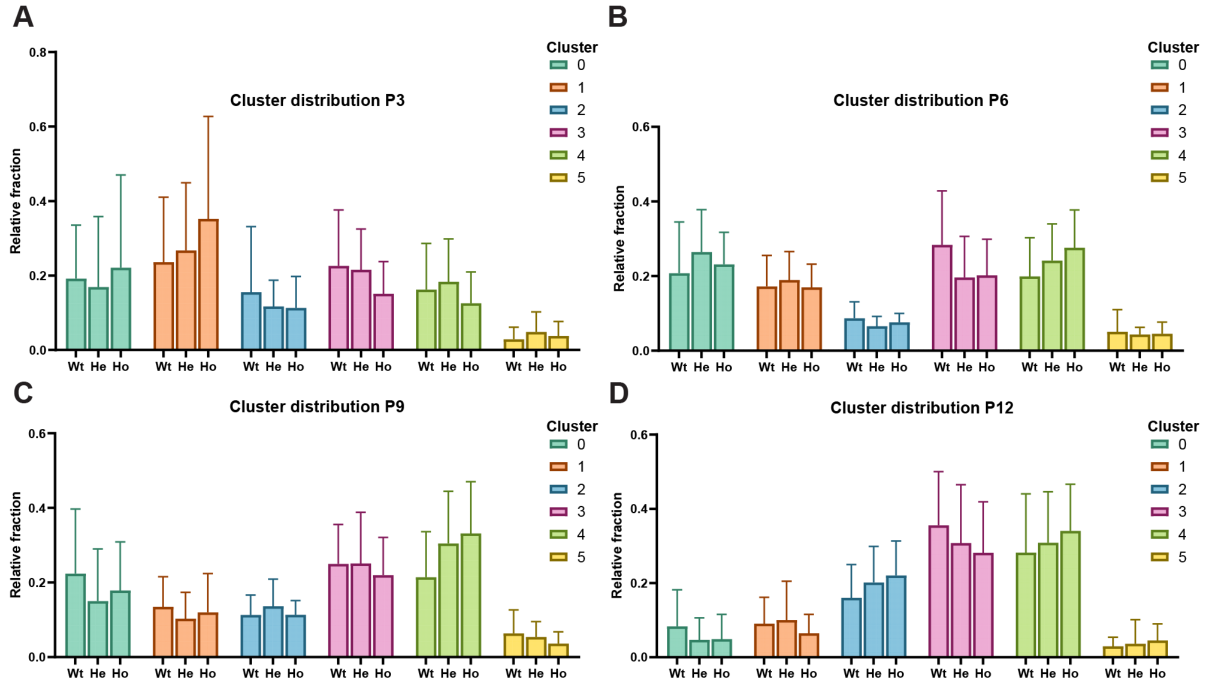


***Supplementary Figure 6: Relative contribution of call clusters in the conditional knockout pups per age and genotype.*** *The relative fraction of calls from wild-type, heterozygous and homozygous conditional knockout pups per cluster at P3 (A), P6 (B), P9 (C) and P12 (D).*

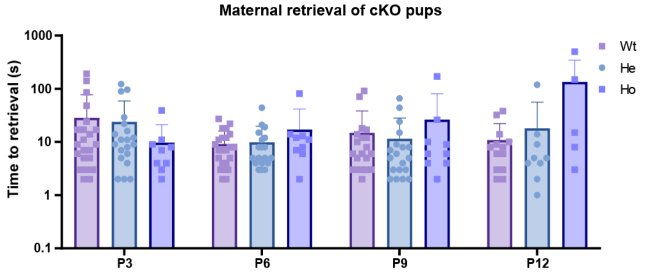


***Supplementary Figure 7: Maternal retrieval time of Pcdh10 conditional knockout pups.*** *Maternal retrieval time of pups over development(P3-P12) and per genotype (Wild-type (Wt), Heterozygous (He), Homozygous (Ho)).*
